# A survey of CIN measures across mechanistic models

**DOI:** 10.1101/2023.06.15.544840

**Authors:** Andrew R. Lynch, Shermineh Bradford, Amber S. Zhou, Kim Oxendine, Les Henderson, Vanessa L. Horner, Beth A. Weaver, Mark E. Burkard

## Abstract

Chromosomal instability (CIN) is the persistent reshuffling of cancer karyotypes via chromosome mis-segregation during cell division. In cancer, CIN exists at varying levels that have differential effects on tumor progression. However, mis-segregation rates remain challenging to assess in human cancer despite an array of available measures. We evaluated measures of CIN by comparing quantitative methods using specific, inducible phenotypic CIN models of chromosome bridges, pseudobipolar spindles, multipolar spindles, and polar chromosomes. For each, we measured CIN fixed and timelapse fluorescence microscopy, chromosome spreads, 6-centromere FISH, bulk transcriptomics, and single cell DNA sequencing (scDNAseq). As expected, microscopy of tumor cells in live and fixed samples correlated well (R=0.77; p<0.01) and sensitively detect CIN. Cytogenetics approaches include chromosome spreads and 6-centromere FISH, which also correlate well (R=0.77; p<0.01) but had limited sensitivity for lower rates of CIN. Bulk genomic DNA signatures and bulk transcriptomic scores, CIN70 and HET70, did not detect CIN. By contrast, single-cell DNA sequencing (scDNAseq) detects CIN with high sensitivity, and correlates very well with imaging methods (R=0.83; p<0.01). In summary, single-cell methods such as imaging, cytogenetics, and scDNAseq can measure CIN, with the latter being the most comprehensive method accessible to clinical samples. To facilitate comparison of CIN rates between phenotypes and methods, we propose a standardized unit of CIN: Mis-segregations per Diploid Division (MDD). This systematic analysis of common CIN measures highlights the superiority of single-cell methods and provides guidance for measuring CIN in the clinical setting.

**Significance:** Cancer relies on genomic changes to drive evolution. One type of change, Chromosomal INstability (CIN), promotes plasticity and heterogeneity of chromosome sets via ongoing errors in mitosis. The rate of these errors informs patient prognosis, drug response, and risk of metastasis. However, measuring CIN in patient tissues is challenging, hindering the emergence of CIN rate as a prognostic and predictive clinical biomarker. To advance clinical measures of CIN, we quantitatively tested the relative performance of several CIN measures in tandem using four well-defined, inducible CIN models. This survey revealed poor sensitivity in several common CIN assays and highlights the primacy of single-cell approaches. Further, we propose a standard, normalized unit of CIN, permitting comparison across methods and studies.

## INTRODUCTION

David von Hansemann and Theodor Boveri described chromosomal instability (CIN) and proposed its role in human cancer over 100 years ago (1, 2). Indeed, CIN accelerates tumor evolution, and portends increased metastasis (3, 4), therapeutic resistance (5–7), and worse prognosis (8–10). Importantly, tumors’ CIN levels vary considerably. While increasing CIN levels correlate with features of advanced cancer, the relationship is not linear or simple. Very high CIN levels inhibit tumors in mouse models (11–13). In human cancer, different CIN levels have been variously correlated with survival, reported both as improved (14–18) and impaired (8, 10, 19, 20). This discrepancy may be due in large part to suboptimal CIN measurements and poorly defined categories of CIN, highlighting the need for accurate, quantitative CIN measures that are amenable to clinical use.

Since von Hansemann and Boveri, several methods of quantifying CIN have been used in basic and translational studies. However, these methods vary in accuracy as well as how comprehensively and directly they assess CIN, leading to disagreement among measures (21, 22). Given these discrepancies, it is important to compare and judge the analytic validity of these approaches to optimize CIN measures for human cancer.

Among cell biologists, direct observation of mitotic abnormalities is a common measure of CIN. Fixed and time-lapse microscopy rely on the visual identification of abnormal mitotic phenotypes such as lagging, polar, and bridging chromosomes as well as spindle multipolarity. This direct observation of mitotic defects is considered the ‘gold standard’. However, the identity of the mis-segregated chromosome(s) remains unclear and, in some cases, even direct observation requires assumptions to infer chromosome mis-segregation, such as whether a lagging chromosome segregates to the correct or incorrect daughter cell (5–7, 23). Further, these methods are not readily applicable to human tumors.

Cytogenetic methods can be used to infer CIN by cell-to-cell variation in chromosome numbers or karyotypes. Cycling cells are captured in mitosis for chromosome counts and karyotypes. However, the reliance on mitotic cells may bias the sample, particularly as *de novo* aneuploidy may delay progression through interphase (24–26). By contrast, centromeric fluorescence *in situ* hybridization (cenFISH) identifies abnormal complements of chromosomes in fixed interphase cells without bias for cycling cells. However, cenFISH is limited to a small number of chromosomes and may suffer from other limitations such as sectioning artifacts. Further, any analysis of living cells in a tumor is biased by cellular selection, as dead cells are cleared (7, 27–29).

Genomics promises a comprehensive evaluation of genomic heterogeneity, and a variety of genomic methods have been proposed to assess CIN. Bulk transcriptional data (bRNAseq) assigns CIN scores to tumor samples based on expression of a selected set of genes. The CIN70 expression score (30), meant to reflect the level of CIN, is derived from an inferred level of aneuploidy and correlate with proliferation and structural aneuploidy. Similarly, the HET70 score correlates with high karyotype heterogeneity in the NCI60 cell line panel, and, unlike CIN70, is independent of proliferation (31). Despite these advances, it is unclear if bulk transcriptional scores can discriminate *ongoing CIN* in the tumor from *historical CIN—* chromosome aberrations that previously arose but do not continue. Similarly, bulk DNA sequencing (bDNAseq) CIN signatures are proposed to quantify and identify the initial cause of CIN, by inferring mechanisms that could produce an observed cell-averaged DNA copy number profile. Whether bulk sequencing measures detect ongoing CIN has not been tested.

Single-cell genomic measures of cell-to-cell variation in copy number analysis are less commonly used in clinical samples but have an advantage over bulk methods as they are not averaged across cells, allowing for direct evaluation of cell-to-cell variation (27). Low-coverage single cell DNA sequencing (scDNAseq) can determine numerical copy numbers of all chromosomes in single cells in experimental models and patient tumors (32, 33). Like FISH, this information can be used to measure CIN by quantifying cell-to-cell heterogeneity in genomics-inferred karyotypes. scDNAseq can be further improved by accounting for cellular selection against highly aneuploid cells using computational modeling and applied Bayesian computation (ABC) (27).

Here, we directly compare these various measures of CIN in well-defined inducible cell-based models. The results reveal the pre-eminence of single-cell measures (microscopic analysis of mitosis, cytogenetic methods, scDNAseq) in measuring ongoing CIN and the inability of bulk molecular sequencing (transcriptional profiles and bulk DNA sequencing) to detect CIN. Among single-cell measures, those that survey all chromosomes across hundreds of cells are most sensitive. The data collected here are made available to investigators who seek to accurately quantify CIN and are an important touchstone to establish the directness of specific measures for ongoing CIN. We anticipate that this comprehensive comparison will allow the most accurate CIN measures to be employed in clinical samples, which will advance mechanistic insight and clinical significance, as well as facilitate use of CIN as a predictive biomarker, such as for microtubule-targeted therapies (17).

## RESULTS

### Design and validation of CIN models with distinct mechanisms

We developed 4 phenotypic models of CIN induced by distinct mechanisms **(Figure 1)**: 1) Bridging chromosomes (Br, red) in CAL51 cells via tetracycline-inducible expression of a dominant negative mutant of telomeric repeat binding factor (*TERF2*-DN) lacking both its basic domain and Myb-binding box (34), 2) pseudobipolar spindles (Pb, blue) in MCF10A cells via induction of centrosome amplification and subsequent spindle pole focusing after tetracycline-inducible expression of polo-like kinase 4 (*PLK4*), 3) multipolar spindles (Mp, green) in MCF10A cells by inhibition of HSET to prevent focusing in the Pb model (17), and 4) polar chromosomes (Po, orange) in MCF10A cells by mitotic enrichment and sequential inhibition of CENP-E then MPS1. This sequential chemical treatment induces anaphase onset with multiple polar chromosomes with high penetrance (35). After induction of each CIN model, we performed imaging (fixed immunofluorescence and 4+ hour time lapse fluorescence microscopy), cytogenetic (mitotic chromosome counts and cenFISH), and sequence-based assays (bRNAseq, bDNAseq, scDNAseq) to measure CIN.

**Figure 1.**
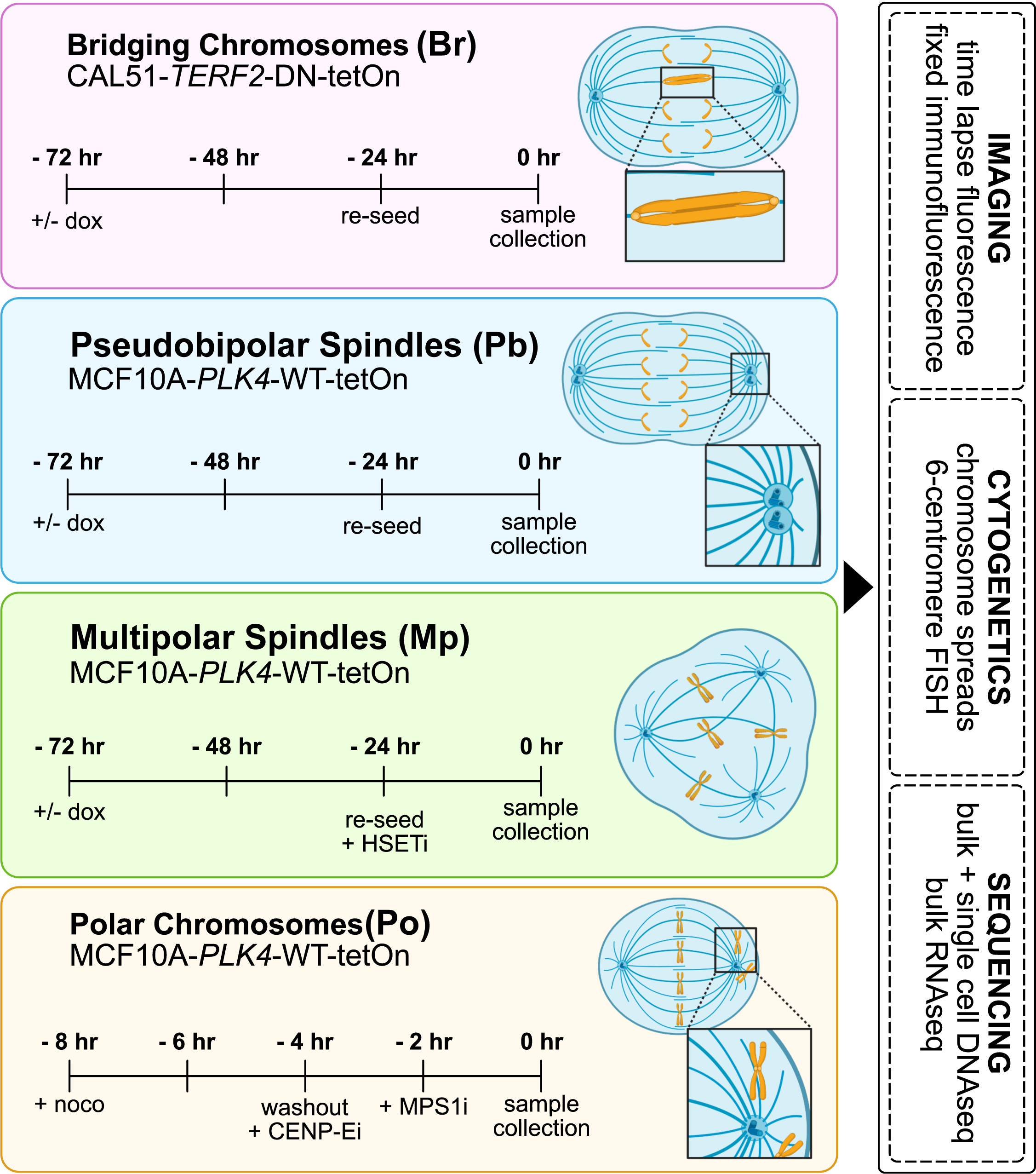
Generation and multimodal analysis of specific inducible CIN phenotypes. Schematic showing genetically and chemically inducible models of CIN phenotypes and subsequent CIN measures employed. Bridging chromosomes (Br) are produced by tetracycline-controlled expression of a dominant negative TERF2 mutant in CAL51 cells. Pseudobipolar spindles (Pb) are produced by tetracycline-controlled expression of WT PLK4 in MCF10A cells. Multipolar spindles (Mp) are produced by further perturbation of Plk4 inducible cells with a chemical inhibitor of HSET (25 μM CW-069). Polar chromosomes (Po) are produced by enrichment of mitotic cells with nocodazole (1 μg/mL), washout into a CENP-E inhibitor (1.5 μM GSK923295) followed by an inhibitor of MPS1 (2 μM AZ3146). All tetracycline-controlled systems were constitutively activated for 72 hours (2 μg/mL doxycycline) before harvest. Po cells used for mitotic chromosome spreads and centromeric cenFISH were washed out at T-0 and harvested 24 hours later after a 2 hour 50 ng/mL colcemid treatment.

### Visualization of mitosis sensitively detects CIN

We first verified that our models induced CIN by microscopy. Using fixed and time lapse imaging, we found the expected CIN phenotypes for each model, indicating successful induction (**Figure 2**). As expected, the Br model induced anaphase bridges in ∼30-40% cells after doxycycline, visible on fixed and live analysis (**Figure 2A,E)**. For the Pb model, we often observed multipolar spindles that focus to pseudobipolar spindles by anaphase onset such that a single spindle pole often has two pericentrin foci (**Figure 2B**). The Pb model displayed pseudobipolar anaphases in 40-50% of cells **(Figure 2B,F)**, though about 5-15% of cells exhibited multipolar anaphase (**Figure 2C,G**). The Mp model induced ∼25% multipolar anaphases after doxycycline + CW-069. The Po model induced high penetrance of CIN with polar chromosomes found in virtually all cells **(Figure 2D-H)**.

**Figure 2.**
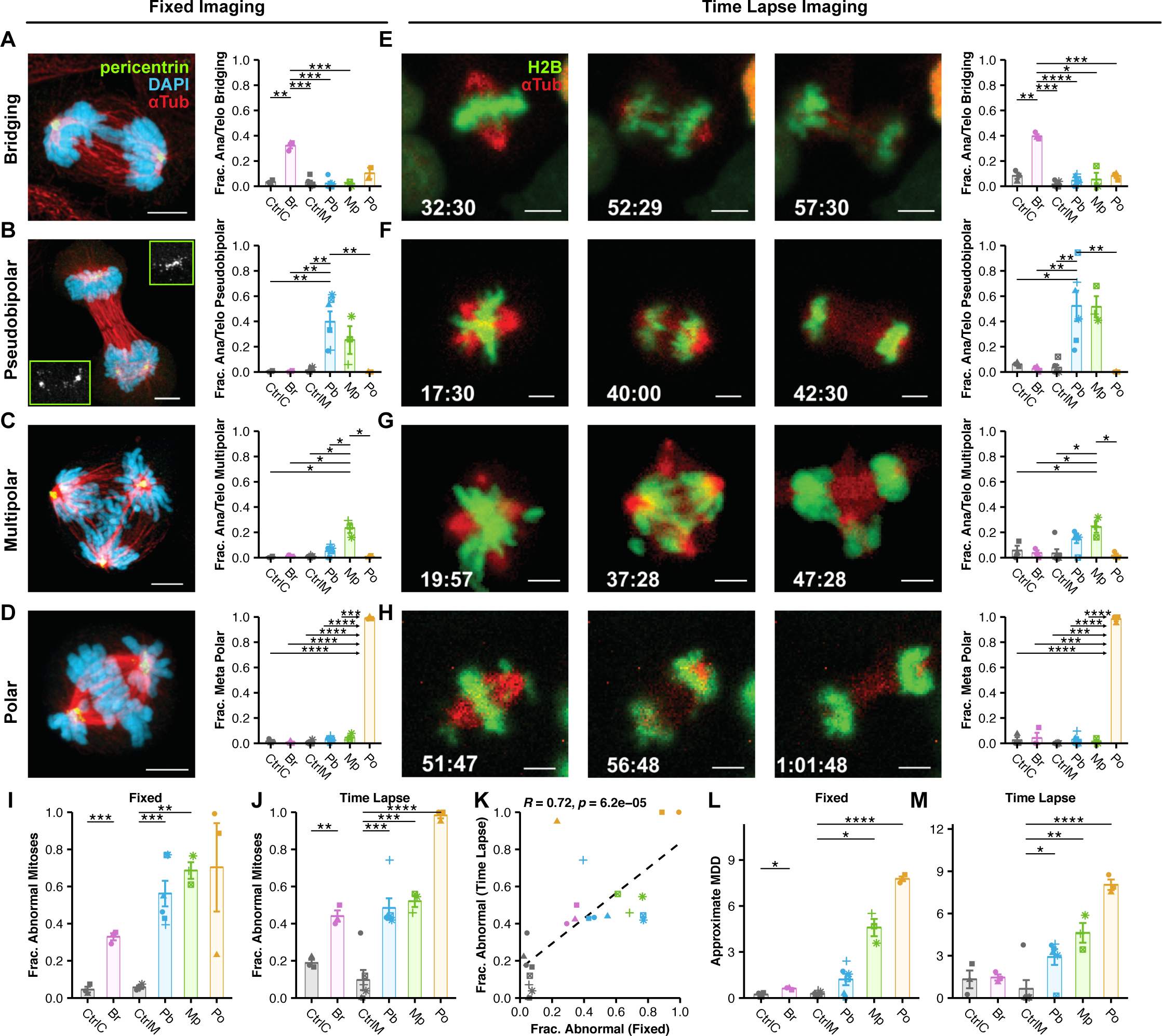
Incidence and specificity of CIN phenotypes via imaging. **(A-D)** Representative images of specific CIN phenotypes in their respective cell line model and quantifications of their observed incidence using fixed immunofluorescence or **(E-H)** time lapse fluorescence imaging. **(I)** Quantification of the fraction of mitoses with any abnormality across phenotypic CIN models for fixed immunofluorescence or **(J)** time lapse fluorescence imaging. Symbolic significance thresholds are 0.05 (*), 0.01 (**), 0.001 (***), and 0.0001 (****). **(K)** Correlation between fixed immunofluorescence and fluorescence time lapse imaging with Pearson correlation coefficient. **(L)** MDD values approximated from fixed imaging. **(M)** MDD values approximated from time lapse fluorescence imaging. MDD values were approximated as the sum of phenotype specific MDD values for each condition **(Supplemental Table 1)**. Colors of individual points indicate the biological replicate. Bars and error bars indicate mean and standard error. Significance values above data are from a two-tailed, two-sample Student’s *t*-tests. For time lapse imaging, N ≥ 20 cells for each of ≥ 3 biological replicates. (Exceptions are single replicates of CtrlC and Pb. See Materials and Methods.) For fixed imaging, N ≥ 50 metaphase and ≥ anaphase/telophase cells for each of ≥ biological replicates. (Exceptions are single replicates of CtrlM, Pb, and Po. See Materials and Methods.)

To evaluate measures of CIN, we quantified the total mitotic aberrations by microscopy using fixed and time lapse imaging **(Figure 2I,J)**. As expected, all had statistically elevated CIN over controls and the two methods were closely correlated **(Figure 2K)**. To directly compare the relative levels of CIN imparted by each model, we quantified a standardized measure of CIN, mis-segregations per diploid division (**MDD**), which we previously described (36). This measure accounts for the relative mis-segregation rates produced by different CIN phenotypes, the relative penetrance of these phenotypes, and the ploidy of the cell. To determine MDD from mitotic observations, we must directly identify mitotic errors. For simplicity, we assume each bridging and lagging chromosomes each produce 1 mis-segregation, polar chromosomes produce 7.8 mis-segregations based on quantitative immunofluorescence imaging of polar centromere foci **(Supplemental Figure 1A)** and multipolar spindles produce 18 mis-segregations based on Lynch et al. 2022 **(Supplemental Table 1, Materials & Methods)**. By this approximation, the mis-segregation rates (MDD) observed for each model are 0.22 ± 0.1 for CtrlC, 0.61 ± 0.04 for Br, 0.28 ± 0.06 for CtrlM, 1.21 ± 0.37 for Pb, 4.59 ± 0.56 for Mp, and 7.77 ± 0.15 for Po by fixed imaging **(Figure 2L)**. By time lapse imaging, these approximations are 1.32 ± 0.63 for CtrlC, 1.45 ± 0.22 for Br, 0.65 ± 0.62 for CtrlM, 2.91 ± 0.56 for Pb, 4.63 ± 0.7 for Mp, and 8.04 ± 0.37 for Po **(Figure 2M)**.

In summary, these findings validate our CIN models and demonstrate low levels of CIN with Br and Pb, intermediate levels with Mp, and high levels with Po. These distinct models and mechanisms of CIN confirm they are suitable models to compare quantitative measures of CIN.

### Short-lived CIN phenotypes are underestimated in fixed imaging

While the two imaging methods correlated well **(Figure 2K)**, time lapse imaging appeared more sensitive to certain CIN phenotypes (**Supplemental Figure 1B)**. For example, we detected significantly more multipolar metaphases and anaphases with lagging chromosomes using time lapse imaging. Anaphase polar chromosomes were more readily detected using time lapse imaging as well. These differentially detected defects are transient in nature, suggesting that the differences are not an artifact of live imaging. For example, multipolar spindles can be clustered rapidly throughout mitosis while lagging and polar chromosomes are often incorporated into the segregating chromosome masses during anaphase. Together, these results indicate that measurement of CIN using fixed imaging, as is common in retrospective clinical analyses of CIN, may underestimate the incidence of some mitotic defects.

### Cytogenetic methods detect CIN with modest sensitivity

Mitotic chromosome counts and centromeric FISH are commonly used cytogenetic approaches to measure CIN (17, 37–39). Chromosome counts detected variation around the modal chromosome count (46 for CAL51s and 47 for MCF10As) in all CIN models and their controls (**Figure 3A**). In several models, in addition to near-diploid aneuploidy there were small fractions of counts consistent with triploid (n=69) or tetraploid (n=92) cells. The variation in chromosome counts only significantly differs from control for the highest rate of CIN generated, the Po condition, in which nearly all cells were affected (**Figure 3B**). Mp divisions may not be detected by this method if these cells delay entry into the next mitosis. cenFISH probe counts can estimate CIN as non-modal counts (**Figure 3C-D**). In some cases CIN measures were similar to controls, likely reflecting the difficulty of measuring CIN by sampling only 6 chromosomes after only 1-2 aberrant divisions. However, nonmodal probe counts were clearly elevated for the models with the highest CIN: two-fold for Mp and three-fold for Po (**Figure 3D**). Recent data suggest a bias for mis-segregation of larger chromosomes, though this can vary based on the insult that causes chromosome missegregation (21, 22, 40). If there is a bias, it may be important to survey chromosomes more broadly than readily feasible with cenFISH. These data suggest that cytogenetics methods are less sensitive than inspection of mitosis at detecting ongoing CIN.

**Figure 3.**
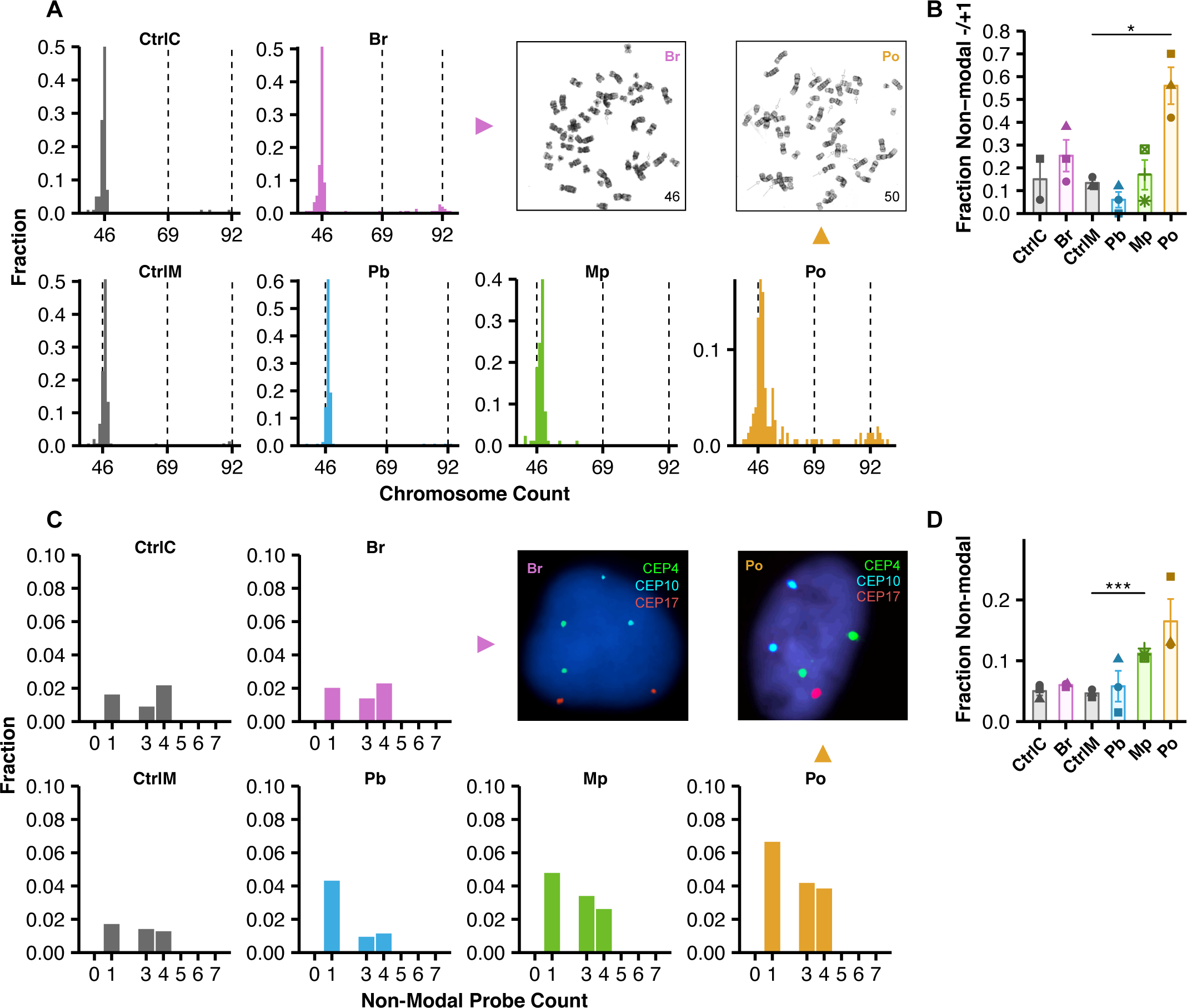
Cytogenetics methods exhibit low sensitivity to ongoing CIN. **(A)** Representative images of mitotic chromosome spreads from each phenotypic CIN model with corresponding histograms of mitotic chromosome counts per spread. **(B)** Quantification of the fraction of mitotic chromosome spreads whose chromosome counts differ from the mode +/- 1. N ≥ 50 mitotic spreads for each of 3 biological replicates. (Exceptions being CtrlC, with 2 biological replicates, and Mp model, which had a low mitotic index in all 3 replicates (see Materials and Methods).) **(C)** Representative centromeric FISH images from each phenotypic CIN model with corresponding histograms of centromeric probe counts per cell. Diploid counts are excluded from histograms to emphasize aneuploid counts. **(D)** Quantification of the fraction of cells whose cenFISH probe counts differ from the mode. Shapes of individual points indicate the biological replicate. N ≥ 200 cells for each of 3 biological replicates. Bars and error bars indicate mean and standard error. Significance values above data are from a two-tailed, two-sample Student’s *t*-tests. Symbolic significance thresholds are 0.05 (*).

### Bulk transcriptomic and genomic CIN signatures do not reflect ongoing CIN

Transcriptional signatures of CIN such as CIN70 and HET70 are used as proxy measures to assess CIN from bulk transcriptomic data in tumor samples (41, 42). These are derived indirectly by identifying gene expression that correlates with aneuploidy in tumor samples (CIN70)(30) and karyotype heterogeneity in cell lines (HET70)(41). CIN70 is partly confounded by proliferation-associated transcription, justifying the development of HET70. Nevertheless, neither CIN70 nor HET70, to our knowledge, have been tested in inducible models of CIN. To determine if these directly measure induced CIN, we employed these transcriptional measures with our models of ongoing CIN. We performed bulk RNA sequencing and measured CIN70 and HET70 signatures in our models **(Supplemental Table 2)**. To validate our results, we verified that doxycycline addition caused a 4-fold increase of *TERF2* expression in the Br condition and a 32-fold increase in *PLK4* expression in Pb and MP. In each case, these were among the top differentially expressed genes **(Supplemental Figure 2A,B)**. Turning to CIN scores, we plotted the distribution of all 70 genes **(Figure 4A,B)** with the mean representing the score. As illustrated, neither CIN70 nor HET70 was increased in any of the CIN models. In fact, CIN70 decreased slightly in Pb and Mp models **(Figure 4A)**, likely due to a decrease in cell proliferation after centrosome amplification (43). This suggests that these bulk transcriptional scores do not detect ongoing CIN.

**Figure 4.**
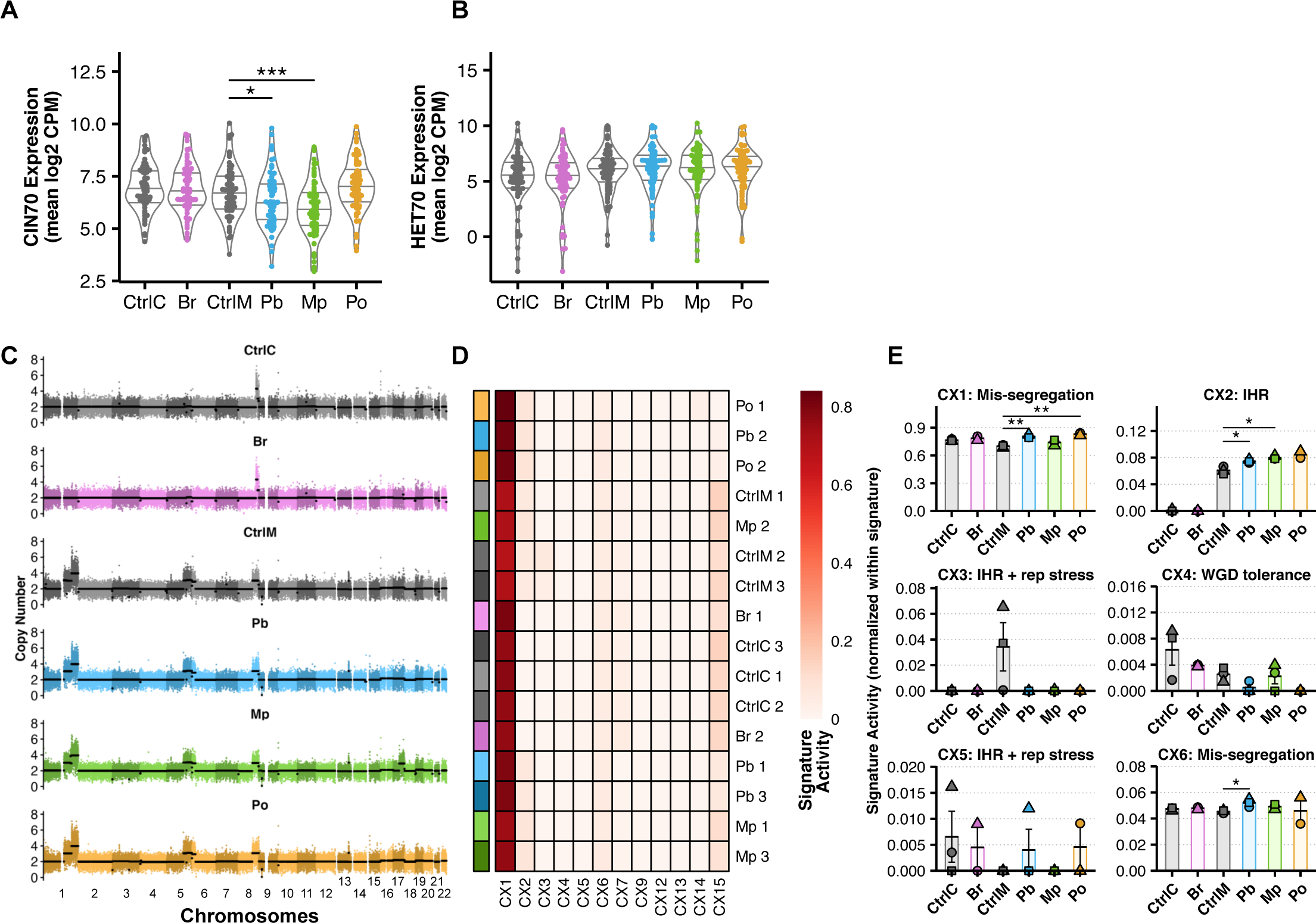
Previously published bulk transcriptomic and genomic CIN signatures do not reflect ongoing CIN. **(A)** Expression levels (mean log2 counts per million (CPM)) of CIN70 and **(B)** HET70 genes from bRNAseq data from each phenotypic CIN model. Each point indicates the average expression level across 3 biological replicates. **(C)** Representative whole-genome copy number profiles of 500 cells from each phenotypic CIN model sequenced together to ∼10x depth. Points indicate copy number of 30 Kb bins. Black lines indicate segment copy number. Alternating shades indicate chromosomes. **(D)** Normalized CIN signature activities of all putative signatures defined in Drews et al. 2022 which provided signal for at least one sample. Color annotations on left of heatmap indicate models and shades indicate replicates. Rows are clustered by similarity of CIN signatures. **(E)** Normalized signature activities of all putative CIN signatures whose mechanistic etiologies are classified as ‘high confidence’ in Drews et al. 2022. CIN signatures for a given sample are normalized to sum to 1. Shapes of individual points indicate the biological replicate. Bars and error bars indicate mean and standard error. Significance values above data are from a two-tailed, two-sample Student’s *t*-tests.

In addition to RNA, bulk genomic DNA measures of CIN are proposed to detect characteristic signatures of copy number variation (CNV) from SNP array and genome sequencing data—essentially measuring patterns of aneuploidy (44). These measures characterize the final state of the tumor, which could either arise from an early event in oncogenesis, or through continuous CIN with selection for certain aneuploid clones. In the latter circumstance, bulk DNA would potentially measure CIN. Whole-genome sequencing (∼10x coverage) and copy number calling in our models revealed nearly identical copy number profiles between control and CIN-induced groups **(Figure 4C)**. We next analyzed recently published CIN signatures (44) in each replicate and clustered the models and controls by signature **(Figure 4D)**. CX1 is the predominant CIN signature in all groups. Importantly, it also appeared in control cells. CX1 corresponds to large scale copy number alterations consistent with whole chromosome or chromosome arm mis-segregation, though is only slightly higher in Pb and Po models than in control cells. Surprisingly, CX6 did not differ between controls and induced CIN models, other than Pb, even though it similarly represents whole chromosome and chromosome arm mis-segregation. CX2, which corresponds to impaired homologous recombination (IHR), is modestly elevated with in Pb, Mp, and Po models, though it is also elevated in control MCF10a cells. **(Figure 4E)**. CX3 was only elevated in MCF10a control cells. Since our models of CIN are induced over 8-72h, they do not provide sufficient time for extensive cellular selection, a process that is likely required for these signatures to appear as they are based on the averaged CNVs of the population. Nevertheless, we conclude that the DNA genomic signatures of CIN do not directly measure ongoing CIN of samples. In sum, we find that bulk genomic measures of CIN, whether transcriptomic or genomic, do not directly measure ongoing CIN, even in an ideal context where tumor purity is not at issue.

### scDNAseq detects ongoing numerical CIN and enables inference of mis-segregation rates

The analyses above suggest that optimal CIN measures would (i) detect all chromosomes, (ii) directly detect CNV variation across cells, and (iii) have high throughput. Single cell DNA sequencing (scDNAseq) meets these characteristics. Even low coverage of reads across the genomes are sufficient to infer copy numbers across all chromosomes. Thus, scDNAseq has been employed to measure cell-cell variation in chromosome copy number and to infer CIN (27, 45). We therefore evaluated scDNAseq as a sensitive measure of CIN in our models.

To evaluate scDNAseq, we sampled 32 single cells per replicate and included a bulk sample of 500 cells to infer average karyotype. **(Supplemental Figure 3)**. We filtered for quality and inferred large-scale chromosome copy number alterations at 1Mb resolution, resulting in 378 high-quality single-cell copy number profiles **(Figure 5A)**. Bulk analyses reveal CAL51 as diploid with a focal 8q amplification; MCF10A cells also had 8q plus gains of 1q, 5q, and recurring sub-clonal gains of Xq. To quantify chromosomal deviations, we evaluated the absolute difference between each single-cell karyotype and the modal karyotype inferred from bulk samples. Both controls—CAL51 and MCF10A—had a small number of deviations from modal karyotype (**Figure 5B**). We did not detect a clear increase in chromosomal deviations in the Br model. This could be due to 1) resolution of chromatin bridges, 2) chromosome breakages resulting in structural variation, but conserved copy number, which would be undetectable by low coverage scDNAseq, or 3) segmental copy number changes smaller than 50% of the chromosome. However, we ruled out the latter possibility using an alternative analysis to detect telomere proximal breaks resulting from breakage of bridging chromosomes, which produced no significant results **(Supplemental Figure 4)**. We found a modest increase in non-modal chromosomes with the Pb model and significant increases in the Mp and Po models, which average about 5-6 deviations per cell respectively **(Figure 5B)**. The percentage of cells with deviations from the mode followed a similar trend, though also detects a small increase in the Br model that is not statistically significance **(Figure 5C)**. As expected, the Mp model displayed a significant bias of chromosome losses over gains **(Figure 5D**)—this is expected since division of duplicated chromosomes into 3+ daughter cells reduces chromosome number. Other models showed no such bias with gains/losses being roughly equal. Taken as a whole, these data support scDNAseq as a sensitive method for detecting CIN.

**Figure 5.**
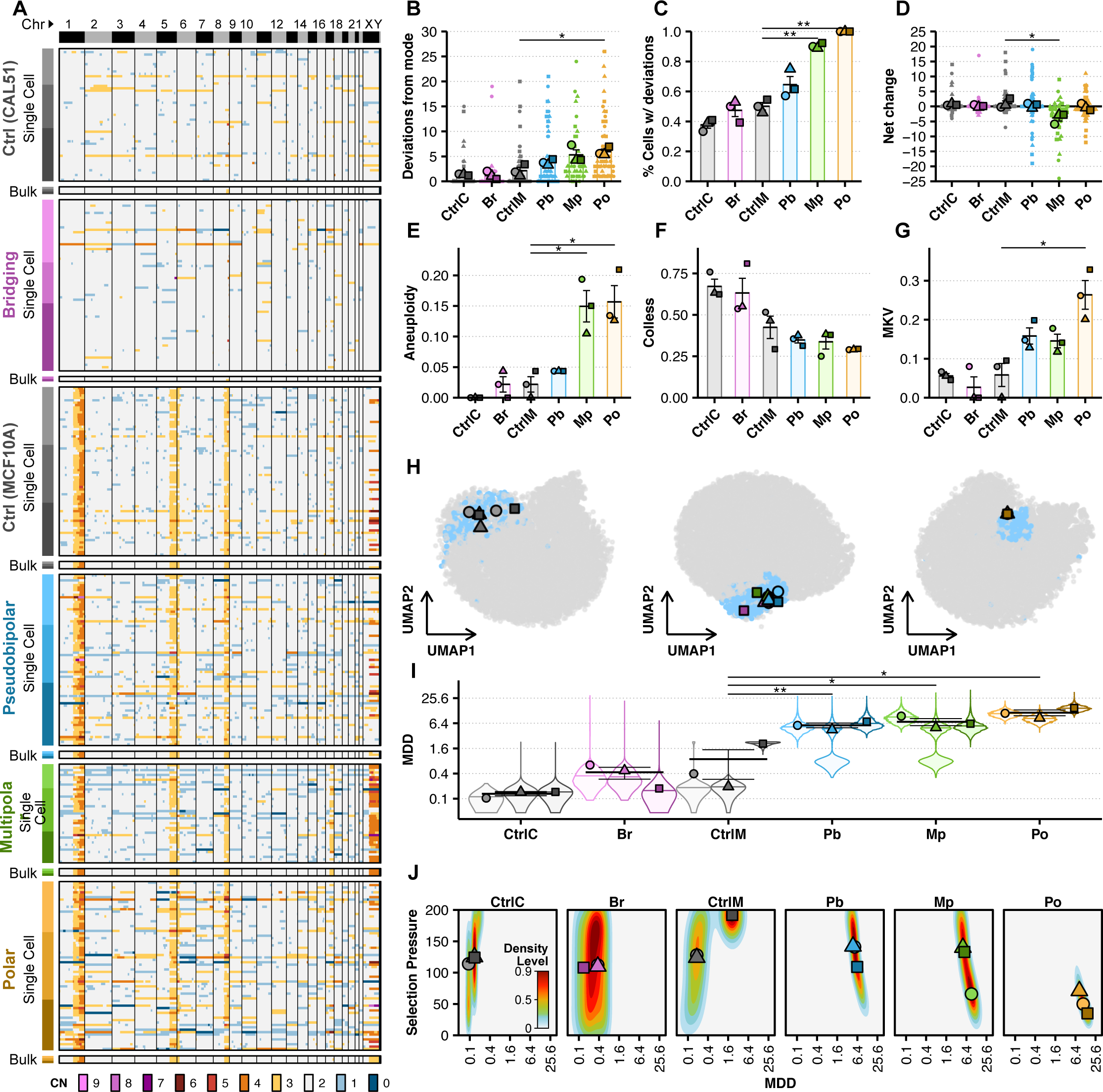
scDNAseq detects ongoing numerical CIN and enables inference of mis-segregation rates. **(A)** Heatmaps of unsorted single cell and bulk (500 cells) copy number profiles low coverage scDNAseq of each phenotypic CIN model. Copy numbers were called using 2.5 Mb genomic bins. Color annotations on left of heatmap indicate models and shades indicate replicates. Color annotations on top of heatmap and vertical lines indicate chromosomes. Heatmap colors indicate copy number. Whole chromosome copy numbers are derived as the mode of bin copy numbers across each chromosome and used to quantify, for each phenotypic model, **(B)** the absolute number of whole chromosome deviations from its modal karyotype, **(C)** the percentage of cells with whole chromosome deviations, and **(D)** the net change of total chromosomes in each cell. Quantified **(E)** aneuploidy (the sub-clonal mean variance within karyotypes), **(F)** Colless indices (i.e., phylogenetic imbalance), and **(G)** mean karyotype variance (MKV, the mean variance of each chromosome) were used as summary statistics for approximate Bayesian computation (ABC). **(H)** Space of summary statistics of independent prior simulation datasets used for ABC projected (small data points) with summary statistics measured in each phenotypic CIN model (large data points) using uniform manifold approximation and projection (UMAP). Blue points indicate accepted simulations for at least one model and replicate, whereas grey were rejected. **(I)** Posterior distributions (violin plots) of mis-segregation rates (mis-segregations per diploid division, MDD) across all replicates for each phenotypic CIN model (individual points) inferred using ABC. A log2 scale is used to better illustrate the data. **(J)** Joint posterior density distributions of accepted mis-segregation rates and selective pressure values from ABC for all replicates of each phenotypic CIN model. Data points are mean values for each replicate. Parameter values for prior simulation datasets are as follows: CtrlC and CtrlM — MDD = [0 … 2.3], S = [0 … 200], Time Steps = [30 … 50]; Br, Pb, and Mp — MDD = [0 … 46], S = [0 … 200], Time Steps = [0 … 4]; Po — MDD = [0 … 46], S = [0 … 200], Time Steps = [0 … 2]. See Materials and Methods for further detail. Shapes of individual points indicate the biological replicate. Bars and error bars indicate mean and standard error. Significance values above data are from a two-tailed, two-sample Student’s *t*-tests.

Single-cell resolution may enable detection of CIN signatures (44), so we repeated our previous analysis using the single cell copy number profiles. Again, the proposed whole/arm mis-segregation signature, CX1, was predominant among the groups **(Supplemental Figure 5)**. However, the relative activity of this signature between groups did not correspond to CIN as observed by microscopy or by directly measuring whole-chromosome copy number alterations. CX6, the other proposed whole/arm mis-segregation signature, had much lower activity among groups and did not correspond to observed CIN. Interestingly, despite their low activity, the only signatures that seemed to reflect the observed trend were decreasing CX4 and increasing CX8, proposed signatures of whole genome doubling and replication stress respectively. This analysis suggests that these measures of CNV at single cell resolution do not well characterize ongoing CIN **(Supplemental Figure 5)**.

Because the fitness levels imparted by different karyotypes can be acted on over time by natural selection, the absolute number of chromosomal deviations in a population may not capture CIN in its entirety. We recently addressed this issue and developed a computational framework to infer mis-segregation rates from scDNAseq datasets using approximate Bayesian computation (ABC), a method to statistically relate biological and simulated data (27). This relies on establishing a phylogenetic tree of single cells and measurements of indexes describing this tree from experimental and simulated data. ABC infers biological parameters from the simulation parameters that produce similar phylogeny.

To employ ABC, we characterized the populations with summary indexes: aneuploidy (average sub-clonal variance within karyotypes), mean karyotype variance (MKV; average variance within chromosomes across the population), and the Colless index, a metric of the imbalance of phylogenetic trees reconstructed from cells’ copy number profiles, an indication of ongoing selection (46, 47) **(Figure 5E-G)**. As expected, our CIN models increased aneuploidy and all but Br increased MKV compared with controls **(Figure 5E,G)**. Further, Colless index decreased indicating a low level of selection is at place after CIN induction **(Figure 5F)**. This likely reflects the lack of selection over the short-time of the experiment (8-72h); by contrast, the control population has been under long-term selection and has higher Colless. Together, the summary statistics enable inference of mis-segregation rates by ABC.

We next applied ABC to infer chromosome mis-segregation rates. First, we performed dimensionality reduction, which verified that the biological data falls within the ‘summary space’ of the simulated data **(Figure 5H)**. Inferring mis-segregation rates (taken as the average of the posterior rate distributions) revealed a wide range across models from 0.3-11.5 MDD (36, 48). There was only a 2-fold increase in mis-segregation rate in the Br model of bridging chromosomes: CtrlC and Br showed mis-segregation rates of 0.12±0.07 and 0.28±0.02 MDD respectively. CtrlM cells had inferred mis-segregation rates at 0.83±0.64 MDD while the Pb, Mp, and Po models had much higher mis-segregation rates of 5.4±0.5, 6.5±1.3, and 11.3±1.8 MDD respectively **(Figure 5I)**. In comparison, the approximated mis-segregation rates of in RPE1 cells is 0.01-0.05 MDD and for U2OS, 0.33-0.46 MDD (36). These rates compare favorably with the number of deviations measured directly from whole chromosome copy number data, particularly when accounting for the partial induction of CIN over 8 hours with the Po model **(Figure 5B)**. They similarly compare with MDD values approximated for each model by imaging methods **(Figure 2L,M)**. Further, joint posterior distributions reflect lower apparent evidence of ongoing selection in the Po model, as compared to the other models, in concordance with the short time-span of CIN induction **(Figure 5J)**. The mis-segregation rate observed in the Mp model, when taking into account the penetrance of the phenotype, agrees with previously observed mis-segregation rates caused by multipolar divisions (27).

We conclude that CIN can be measured several ways using single cell copy number profiles. Copy number alterations can be calculated directly, particularly if the period of CIN is relatively short, in which case karyotype selection is not a significant factor. Inferring chromosome mis-segregation rates by comparing to simulated data works in both short and long time periods, though it is ideal for long time periods when karyotype selection becomes a strong confounder.

### Concordance and performance of CIN measures

To assess and summarize the performance of CIN measures, we performed standardized effect size and pairwise correlation analyses. As expected, fixed immunofluorescence, time lapse, non-modal mitotic chromosome, and centromeric probe counts were significantly correlated (α = 0.01), excepting the pair of fixed imaging and chromosome counts **(Figure 6A)**. By contrast, bulk transcriptional CIN signatures (CIN70, HET70) did not correlate to imaging, nor cytogenetics, nor between themselves. Considering genomic signatures on bulk (CX1, CX6 bDNAseq), and with single cell data (CX1 scDNAseq; CX6 scDNAseq), these also correlated poorly with one another with the exception of single-cell CX1 and CX6. Interestingly, the CIN signature CX1, when measured in bDNAseq data, did modestly correlate with time-lapse imaging, but not fixed imaging. HET70 negatively correlated with the CX1 and CX6 signatures in single cells. This seemed to be a cell line-dependent effect as MCF10A cells had relatively high HET70 expression and low CX1 and CX6 signature activity than CAL51, regardless of CIN model **(Supplemental Figure 6)**. scDNAseq chromosomal analyses performed well whether they considered whole-chromosome deviations from mode or inferred MDD from ABC—not only do these correlate with each other, but also with fixed imaging, time lapse, and FISH analyses. We conclude that the single cell analyses perform well for measuring rates of ongoing CIN.

**Figure 6.**
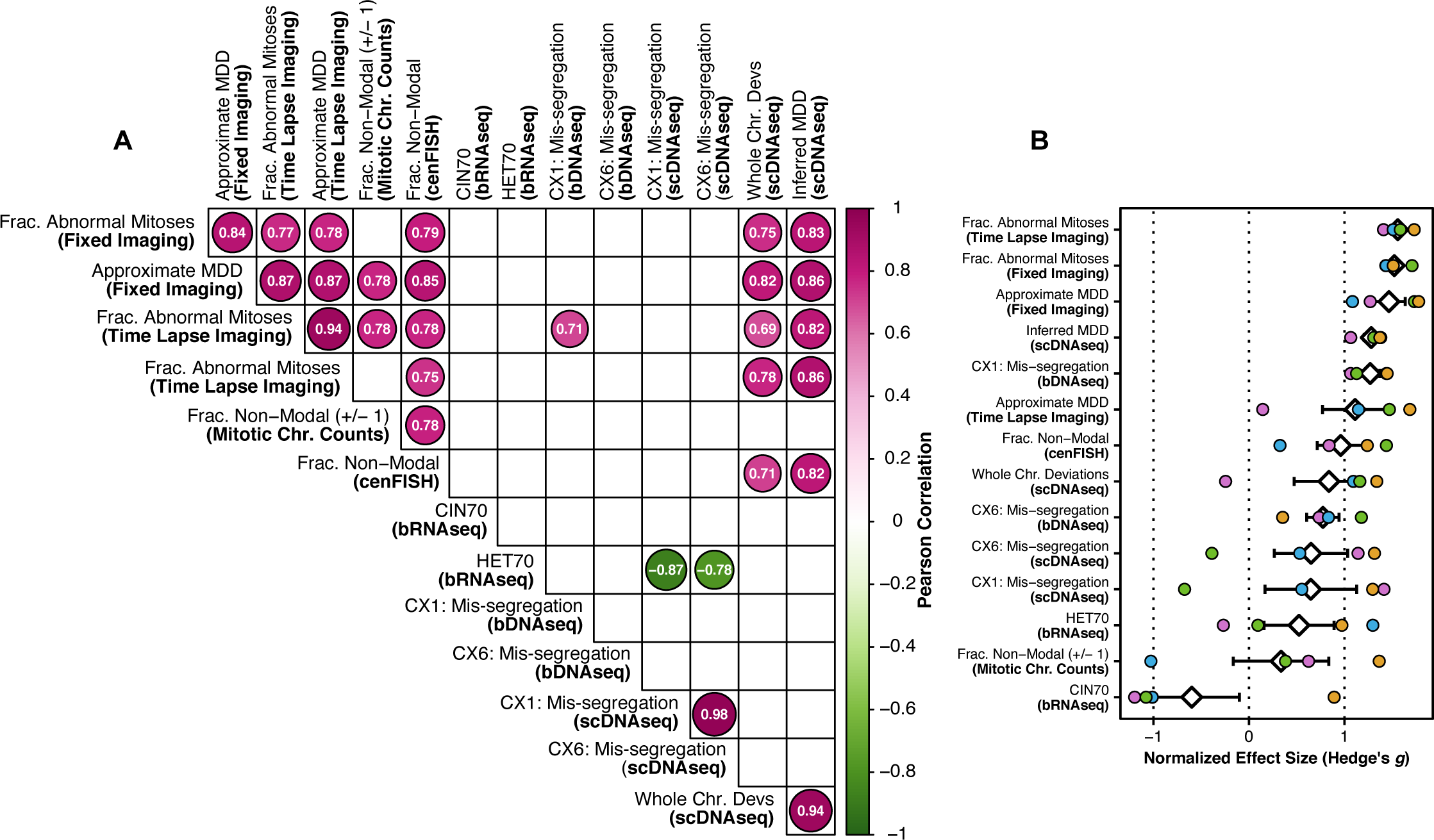
Concordance and performance of CIN measures. (A) Pairwise correlations of CIN measurements. Only statistically significant Pearson correlation coefficients (α = 0.01) are shown. Size of circles represents degree of correlation. (B) Effect sizes (Hedge’s *g*) of CIN measurements for each model compared to its control. White diamond and error bars indicate mean and standard error.

We next turned to effect size, Hedge’s *g*, which allows us to compare sensitivity of different measures (**Figure 6B**). Average standardized effect sizes across all CIN phenotypes were high among the imaging methods. Inferred MDD from scDNAseq was also very high, as expected form sampling chromosomes and accounting for cell selection. Next, CX1 measured by bDNAseq was also high though, as noted previously, these scores do not reflect observed CIN levels by imaging or single-cell sequencing. The effect size for cenFISH was similar to whole chromosome deviations in scDNAseq. The CIN signature of mis-segregation (CX6), as measured by bulk or single cell DNAseq was middling. Transcriptional measures of CIN (CIN70 and HET70) and CIN measured by mitotic chromosome counts exhibited the lowest effect sizes. We conclude that highly sensitive quantitative measures of CIN can be achieved by direct microscopy or by scDNAseq combined with inference of rates by ABC.

In summary, our data compare proposed and widely used measures of CIN on simplified models where CIN is induced. Paradoxically these exhibit vastly different capabilities in detecting CIN. Direct observation of CIN by imaging is the most sensitive and reliable followed by scDNAseq measures and cenFISH. By contrast, transcriptional CIN scores, mitotic chromosome counts, and statistically derived CIN signatures, regardless of bulk or single cell resolution, were far less capable in measuring ongoing CIN in terms of effect size and correlation with other measures.

## DISCUSSION

Ongoing CIN is defined by an elevated rate of chromosome mis-segregation, which varies across tumors and depends on the penetrance and error frequency of specific CIN mechanisms. The intrinsic rate of mis-segregation in a cell population produces functional consequences for tumorigenesis, cancer progression, and treatment response. Very low mis-segregation rates limit tumorigenesis, presumably due to reduced adaptive potential, while high mis-segregation rates have the same effect through loss of necessary genetic material. Intermediate mis-segregation rates can promote tumorigenesis. Once a tumor has formed, reports conflict on whether patients whose tumors exhibit moderate to high intrinsic rates of mis-segregation tend to have poorer clinical outcomes (14–18) (8, 10, 19, 20), likely due to limitations of current methods for measuring CIN in patient tumors. Breast cancer patients whose tumors exhibit high mis-segregation rates tend to have improved response to taxanes, likely because CIN sensitizes to the multipolar mitotic spindles produced by these drugs. Despite the clinical significance of CIN and its variability across tumors, its measurement is not currently used to guide patient care (8, 12, 13, 17, 49).

The reason CIN is not measured for clinical use is ultimately due to three factors: accessibility, scalability, and sensitivity. In reviewing the CIN measures tested in this study, we find varying levels of accessibility and scalability **(Table 1)**. Some measures are clinically accessible in terms of necessary equipment and reagents, but they do not easily scale because of time required for sample preparation or data acquisition. For example, fixed imaging is commonplace for pathological assessment of tumors. In this study, fixed immunofluorescence imaging was among the most sensitive methods to detect ongoing CIN. However, the quantification of CIN phenotypes in these samples is not scalable as data acquisition requires substantial time commitments and sampling a sufficient number of mitoses requires highly proliferative tumors, such as diffuse large B-cell lymphoma (8). Karyotypes and chromosome counts are not readily possible in patient tissues, as they require a large number of mitotic cells, not easily obtained by culturing tumor tissues. Likewise, time lapse imaging is not possible directly in patient tumors, and is laborious, time-consuming, and expensive even with *ex vivo* culture, fluorescent labeling, and imaging of patient organoids.

**Table 1.**
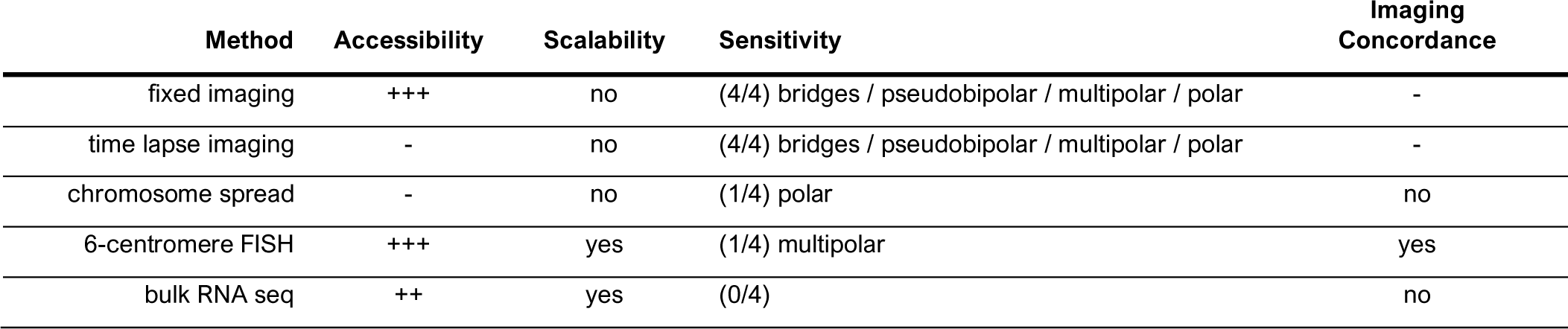
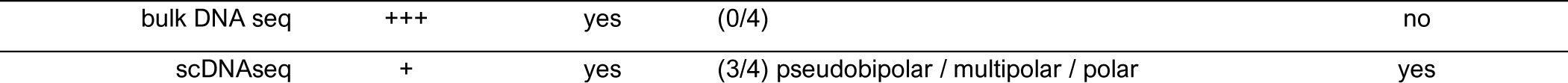
Characteristics of CIN measurement methods. Categories for clinical accessibility are (-) insurmountable barriers, (+) requires special equipment and/or reagents **<u>and</u>** time-consuming and/or laborious, (++) requires special equipment and/or reagents **<u>or</u>** time-consuming and/or laborious, or (+++) commonly performed clinical assays use the same equipment and reagents. Scalability is determined by the pre-existence of established protocols and/or platforms for high throughput sample preparation and data acquisition in a clinical or non-clinical setting. Sensitivity to a specific mechanism is determined by a significant difference between the control and CIN-induced groups. Imaging concordance is determined by significant correlation to both imaging methods.

Measures of CIN based on bulk genomic and transcriptomic sequencing are attractive due to the wide availability of shared high-throughput sequencing data, and because they are already in routine clinical use. However, bulk sequencing failed to detect ongoing CIN in any of our phenotypic models. This is likely because, despite the misleading nomenclature, bulk sequencing measures detect RNA or DNA levels averaged across many cells and thus establish an average aneuploid state, without the ability to detect cell-to-cell differences in chromosome copy numbers. One fundamental problem in the field of genomic integrity is that CIN and aneuploidy are often conflated. To be fair, aneuploidy is a product of prior mis-segregation events, providing a rationale to infer a degree of CIN from the current degree of aneuploidy in a tumor. We can describe this as *historical CIN*, which represents an accumulation of prior chromosome gains/losses and cellular selection. However, this does not permit inference of ongoing chromosome gains or losses because any degree of aneuploidy could, in fact, be a product of a single event that was not repeated. By contrast, *ongoing CIN*, described here, is a process in which there are ongoing aberrations over multiple cell divisions, which is associated with metastasis and response to microtubule-targeted therapies (8, 12, 13, 17, 49).

The *de novo* karyotype heterogeneity in cell populations with ongoing CIN does not significantly alter the predominant karyotype and is not detected by bulk genomic CIN signatures (e.g., CX1 and CX6). This demonstrates that these signatures detect historical aneuploidy rather than ongoing CIN. Nor do the gene expression states of new aneuploid clones significantly alter the predominant transcriptional phenotype with respect to previously established transcriptomic CIN scores (CIN70 and HET70). As for the CIN70 score, this result was unsurprising for two reasons — 1) the derivation of CIN70 was based on the fundamental assumption that aneuploidy is derived by CIN and 2) CIN70 is confounded by upregulation of proliferative genes, as aneuploidy, proliferation, and advanced tumor grade are all co-variates. Further this result comports with previous findings (41, 50). MCF10A cells scored slightly higher on HET70 than CAL51. While this may represent cell line specific gene expression, we note that the scDNAseq data shows MCF10A cells exhibiting higher karyotype heterogeneity than CAL51 at baseline, despite exhibiting similar rates of mitotic errors during imaging. Thus, while HET70 scores did not respond to induced CIN and karyotype heterogeneity, we cannot rule out that HET70 detects a pre-existing transcriptional phenotype that is tolerant of aneuploidy and increased karyotype heterogeneity. However, while HET70 did not detect induced, ongoing CIN in any of our models, MCF10A cells did score higher on HET70 than CAL51. While this may represent cell line specific gene expression, we note that the scDNAseq data shows MCF10A cells exhibiting higher karyotype heterogeneity than CAL51 at baseline, despite exhibiting similar, if somewhat lower rates of mitotic errors during imaging. Although HET70 scores did not increase with induced CIN and karyotype heterogeneity, we cannot rule out that HET70 detects a transcriptional phenotype that correlates with aneuploidy tolerance.

As a clinically accessible alternative to imaging, scDNAseq provides the best measure of CIN in terms of sensitivity and correlation to both fixed and time lapse imaging. CIN can be measured by quantifying the absolute number of chromosomal deviations from the modal (i.e., clonal) karyotype of a population. Additionally, the rate of mis-segregation resulting in a given population of single cell copy number profiles can be measured by pairing computational modeling and statistical inference. Both analysis methods performed well in measures of ongoing induced CIN over relatively few cell divisions. However, we have previously found that the latter, inference of CIN, performs better in the context of longer time scales, such as the growth of a tumor (27). While single-cell sequencing is not currently used in clinical care, major advances in ultra-high-throughput sequencing will likely make low-coverage scDNAseq accessible at clinical capacity (51–53). Toward this end, we estimated that ∼200 cells are needed for accurate measurement of CIN (27). Although we did not detect increased segmental copy number alterations caused by bridging chromosomes in our inducible model, further advancement of sequencing technology and scDNAseq methods may enable more robust detection of copy-neutral structural variation in single cells. Importantly, the measurement of structural CIN using scDNAseq would require relatively uniform coverage of the entire genome, which is not provided by mutation panels. For intact tumors, it would be important to distinguish the genome structures of tumor cells from stromal cells, which could be accomplished by simultaneously detecting cancer-specific mutations.

One limitation of this study is that we measured CIN in cell-line models a short time after inducing CIN. This approach is ideal to ensure ongoing CIN, but we cannot rule out that methods that failed in these models would correlate with ongoing CIN after cellular selection or would indicate a permissive cellular context for ongoing CIN. For example, although we found that cenFISH only detects the highest levels of CIN in our system, we cannot rule out that this method becomes more sensitive over several generations with elevated CIN, which would increase the probability that one of a limited set of probes would detect an alteration. Further, in this study, we did not evaluate single cell RNAseq (scRNAseq) as a method of measuring CIN. Currently, there are no widely used scRNAseq-based measures of CIN and we did not seek new methods of CIN measurement. However, given the preponderance of evidence of the transcriptional consequences of CIN and aneuploidy (26, 54–58), it is conceivable that robust transcriptional CIN signatures could be derived at single cell resolution. Whether these would reflect gene dosage, a general response to CIN or would be phenotype/mechanism-dependent is unclear. In any case, the reliability of these measures could be limited by dosage compensation (59–63). Although we could select specific genes unaffected by compensation, the sparse data with current scRNAseq platforms remains a challenge. On the other hand, large-scale DNA copy number alterations can be inferred from scRNAseq data (64, 65), which would provide an additional dimension of single cell genomic data and perhaps increase the reliability of a measure of CIN based on single cell chromosome copy number data.

This work provides a thorough empirical analysis of the relative capability of current CIN measures to detect ongoing CIN across specific, inducible phenotypic models of CIN. We find current measures differ in their ability to detect ongoing CIN and that some fail to reliably detect CIN at all. Imaging approaches are the most sensitive and reliable. Cytogenetic approaches have low sensitivity, only significantly detecting the model which had among the highest mis-segregation rates. Bulk genomic and transcriptomic measures do not reflect ongoing CIN while single cell genomic methods, particularly the inference of mis-segregation rates, offer both sensitivity and potential for clinical accessibility. In light of these conclusions, we recommend single-cell genomics with Bayesian inference as the best method for further development of a clinically accessible measure of CIN. Importantly, the models and data generated here serve as a resource for investigators seeking to validate innovative measures of ongoing CIN from bulk and single-cell sequencing.

## MATERIALS AND METHODS

### Cell line derivation and cultivation conditions

MCF10A-PLK4-WT-tetOn cells were kindly provided by the laboratory of Dr. David Pellman (66). We used lentiviral transduction to stably express H2B-mNeonGren and mScarlet-α-Tubulin. CAL51 cells were obtained from DSMZ-German Collection of Microorganisms and Cell Cultures and CAL51-TERF2-DN-tetOn was generated using retroviral transduction of pCMV Retro TetO into which we cloned the TERF2-DN mutant sequence (Addgene, 16069) and an mScarlet reporter. To make retrovirus, we transfected 293T cells then transduced CAL51 cells expressing TetR for ∼18 hours with 4 ug/mL polybrene. We selected with puromycin (2 ug/mL) and sub-cloned in 96-wells to generate monoclonal lines. All cell lines were tested and free from mycoplasma contamination prior to study.

Cells were maintained at 37°C and 5% CO_2_ in a humidified, water-jacked incubator and propagated in either Dulbecco’s Modified Eagle’s Medium (DMEM)/High Glucose (Cytiva Hyclone, SH3024301) for CAL51 cells or mammary epithelial basal media for MCF10A cells, consisting of DMEM/F12 (Cytiva Hyclone, SH3026101), 5% horse serum (Gibco, 16050122), 20 ng/mL EGF (Peptrotech, AF-100-15), 0.5 mg/ml hydrocortisone (MP Biomedicals, 0219456901), 100 ng/ml cholera toxin (Enzo Life Sciences, BMLG1170001), and 10 ug/ml insulin (Millipore Sigma, I9278). All growth media is supplemented with 10% fetal bovine serum (GeminiBio, 900-108),100 units/mL penicillin-streptomycin (Gibco, 15070063), and plasmocin prophylactic (Invivogen, ant-mpp) to a final concentration of 5 ug/mL.

### Statistical Analyses

Statistical analysis was completed in R/Rstudio (v4.2.3/v1.2.5019) (67, 67). Unless otherwise specified, statistical significance between group means is determined using two-tailed, two-sample Student’s *t*-tests and α = 0.05 over at least 3 biological replicates.

### Fixed immunofluorescence microscopy

Cells were re-seeded on sterile glass #1.5 cover slips (Fisher, 12-545-81P) at 40% and grown to ∼70-80% over 18-24 hours. Coverslips are rinsed in warmed (37°C) PBS (x1) before fixation in warmed 4% paraformaldehyde (PFA) in PHEM buffer (120 mM PIPES, 50 mM HEPES, 20 mM EGTA, 4 mM MgSO_4_ in pure water, pH 7.0) for 10 minutes. Coverslips are rinsed of fixative (x3), extracted with 1% NP40 in PHEM buffer for 15 minutes, and blocked with 3% bovine surum albumin (BSA)(Millipore Sigma, A2153) and 0.1% Triton X-100 in PBS (BSA + PBSTx) for 30 minutes. Primary and secondary antibodies were pooled separately in BSA + PBSTx. Coverslips were incubated with primary antibodies for 1 hour at room temperature, rinsed (x3) in PBSTx, then incubated with secondary antibodies for 30 minutes, and rinsed (x3) again in PBSTx. Coverslips were counterstained with 10 ug/mL DAPI (Sigma-Aldrich, 62248), mounted on glass slides with Prolong Dimaond anti-fade medium (Molecular Probes, P36970), and cured for 48 hours. Cells were immunostained to visualize α-tubulin (Primary — 1:1000 mouse (γ1) anti-α-tubulin (DM1A)(Invitrogen, 14-4502-82, RRID: AB_1210456; Secondary — 1:350 goat anti-mouse (γ1) + Alexa Fluor 750 (Invitrogen, A-21037, RRID: AB_2535708) for CAL51 cells or goat anti-mouse (H+L) + Alexa Fluor 555 (Invitrogen, A-11001, RRID: AB_2534069) for MCF10A cells) and pericentrin (Primary — 1:1000 rabbit anti-pericentrin (Abcam, ab4448); Secondary — 1:350 chicken anti-rabbit (H+L) + Alexa Fluor 647 (Invitrogen, A-21443, RRID: AB_2535861) for CAL51 cells or goat anti-rabbit (H+L) + Alexa Fluor 488 (Invitrogen, A-11008, RRID: AB_143165) for MCF10A cells)

Image acquisition was performed on a Nikon Eclipse Ti inverted microscope equipped with motorized stage, LED epifluorescence light source (Spectra X), 60x/1.4NA (Plan Apo) DIC oil immersion objective, and ORCA Flash4.0 V2+ digital sCMOS camera (Hamamatsu). Optical sections were taken at 200-nm intervals and deconvolved using the LIM 3D Deconvolution module in Nikon Elements. Micrograph montages were prepared in Affinity Designer.

For analysis, mitotic phenotypes of all mitotic cells observed in metaphase, anaphase, or telophase were recorded until at least 50 metaphase cells were counted or until all four coverslips were expended. Exceptions are replicate 2 of Po, with 10 metaphase cells and 137 anaphase and telophase cells, replicate 3 of CtrlM with 30 metaphase cells and 21 anaphase and telophase cells, and replicate 3 of Pb, with 11 metaphase cells and 3 anaphase and telophase cells. Data from these replicates were retained and reported as the alternative measures of the replicate provided sufficient information.

### Time lapse fluorescence microscopy

Cells were re-seeded in 4- or 8-well chamber slides (Ibidi, 80426) at 40% and grown to ∼70-80% over 18-24 hours. At the time of re-plating, CAL51-TERF2-DN-mCherry-TetOn cells were transduced with adenovirus to express β-tubulin-mScarlet + P2A-H2B-NeonGreen to visualize mitotic spindles and DNA. They were then imaged on a Nikon Ti-E2 inverted fluorescence microscope outfitted with a Yokogawa CSU-W1 spinning disk confocal scanning unit. Images were collected every 2.5 minutes for 4-12 hours with a 20x/0.5NA (P Fluotar) objective and an ORCA Flash 4 digital sCMOS camera (Hamamatsu). Environmental control was maintained by a humidified, stage-top chamber (Tokei Hit) set to 37°C and 5% CO_2_. Micrograph montages were prepared in Affinity Designer.

For analysis, at least 20 cells for each replicate and each CIN model were tracked from nuclear envelope breakdown through to cytokinesis and mitotic phenotypes were recorded for metaphase, anaphase, and telophase. Exceptions are replicate 1 of CtrlC, with 17 cells, and replicate 5 of Pb, with 18 cells. Data from these replicates were retained and reported as the alternative measures of the replicate provided sufficient information.

### Imaging-based approximation of mis-segregations per diploid division (MDD)

MDD was approximated using fixed immunofluorescence and time lapse fluorescence imaging data as well as the formula 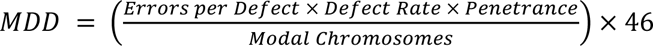, as previously described (36). To derive the values for this formula, we made the following assumptions: 1) Lagging and bridging chromosomes are assumed to produce 1 mis-segregation; 2) Polar chromosomes are assumed to produce 7.8 mis-segregations as we observed cells treated under the Po model (prior to anaphase onset) had an average of 15.6 polar ACA foci **(Supplemental Figure 1A)**; 3) Pseudobipolar spindles do not produce mis-segregation events; 4) Multipolar spindles are assumed to produce 18 mis-segregations based on Lynch et al. 2022; 5) CAL51 cells have 46 chromosomes and MCF10A cells have 47 chromosomes; 6) The penetrance of each observed CIN phenotype is 100% (i.e., 100% chance the defect results in mis-segregation). The fraction of cells with polar chromosomes represents metaphase cells wherein they are most readily detectable. All other CIN phenotypes are taken from anaphase or telophase cells.

### Cytogenetics

Cells were re-seeded in T75 flasks at 40% and grown to ∼70-80% density over 18-24 hours prior to harvest. For Po, AZ3146 was washed out at T-0 hours and harvest performed at T+24 hours. Colcemid (Millipore Sigma, 234109) was added to a final concentration of 50 ng/mL and incubated for 2 hours to enrich mitotic cells. We retained the media and trypsinized cells, rinsing once in warm PBS. Cells were resuspended and swelled for 5 minutes in 75 mM KCl with 10 drops of Carnoy fixative (3:1 methanol:glacial acetic acid). This was followed by three successive resuspensions in Carnoy fixative, after which, samples were stored at −30°C then cells were dropped onto slides and dried in a drying chamber. For analysis, at least 50 mitotic chromosome spreads were counted for each replicate of each model with the exception of the Mp model, which exhibited a relatively few mitotic cells in each biological replicate (18, 32, and 35 mitotic chromosome spreads).

Fluorescence in situ hybridization (FISH) enumeration of chromosomes 3, 4, 7, 9, 10, and 17 was completed using the following 2 probe mixes: Vysis CEP 3 (D3Z1) labeled SpectrumOrange (Vysis, 06J3613) localizing to 3p11.1-q11.1, Vysis CEP 7 (D7Z1) labeled SpectrumAqua (Vysis, 06J5427) localizing to 7p11.1-q11.1, Vysis CEP 9 labeled SpectrumGreen (Vysis, 06J3719) localizing to 9p11-q11 in IntelliFISH hybridization buffer (Vysis, 08N8701), and Vysis CEP 4 labeled SpectrumGreen (Vysis, 06J3714) localizing to 4p11-q11, Vysis CEP 10 labeled SpectrumAqua (Vysis, 06J5420) localizing to 10p11.1-q11.1 and Vysis CEP 17 (D17Z1) labeled SpectrumOrange (Vysis, 06J3697) localizing to 17p11.1-q11.1 in IntelliFISH hybridization buffer (Abbott Molecular, Des Plaines, IL). Slides were aged for 2 minutes in 2xSSC at 74°C and put through a dehydration ethanol series (70%, 85%, and 95%). Slides and probe were codenatured by heating at 80°C for 2 minutes using a ThermoBrite instrument (Abbott Molecular). Hybridization was performed overnight at 37°C. Finally, the slides were mounted with Vectashield containing DAPI (Vector Laboratories). Localization of the probes was confirmed on pooled cytogenetically normal blood controls. For analysis, at least 100 interphase cells were counted per 3-probe set.

### Single-cell DNA sequencing and analysis

### Single cell DNA sample preparation

Cells were re-seeded in 6-well plates at 40% and grown to ∼70-80% density over 18-24 hours prior to harvest. Cells were washed with HBSS, trypsinized, pelleted, and washed by resuspension in wash buffer (2% FBS in 1x PBS). Pelleting once more, cells were resuspended in 500 µL wash buffer and fixed by dropwise addition to 4.5 mL 80% ethanol. Samples were stored at −30°C.

### Flow cytometry and fluorescence activated cell sorting

Fixed cell suspensions were pelleted and resuspended in wash buffer containing 10 µg/mL DAPI. Single cells or bulk samples (500 cells) were sorted by FACS (BD FACS AriaII), gating on 0.5-1.5x the DAPI signal intensity of the G1 peak, through a 130 μm low-pressure deposition nozzle into 96 well PCR plates containing 10 µL 1x prepared Cell Lysis and Fragmentation Buffer Mix (Millipore Sigma, WGA4). Doublets were excluded from analysis via standard FSC/SSC gating procedures. DNA content was analyzed via DAPI excitation at 355 nm and 450/50 emission using a 410 nm long pass dichroic filter. Gating for cell sorting was defined by 0.5x/1.5x (lower/upper) the DAPI intensity of the G1 peak. After sorting, the PCR plates were immediately centrifuged at 100 x g for 1 minute before library preparation. Plates were kept at 4°C when not on the cell sorter.

### Single cell DNA library preparation

Initial cell lysis, genomic fragmentation, and genomic amplification reactions were done with the GenomePlex® Single Cell Whole Genome Amplification Kit (Millipore Sigma, WGA4). Initial genomic library distributions were assessed on a 1.5% agarose gel and purified using a ZR-96 Genomic DNA Clean & Concentrator-5 Kit (Zymo Research, D4067). Library concentrations were quantified using the Quant-iT™ dsDNA Broad Range Assay Kit (Invitrogen, Q33130) and normalized before additional preparation. Genomic libraries were enzymatically fragmented to ∼250 bp, 5’-phosphorylated, 3’-dA-tailed, and index adaptor-ligated with the sparQ DNA Frag & Library Prep Kit (Quantabio, 95194). Ligated adaptors were standard P5 and custom uniquely indexed P7 illumina adaptors described previously (68). Indexed libraries underwent eight additional amplification cycles, purification with Axygen® AxyPrep MAG PCR Clean-Up beads (Corning, MAG-PCR-CL), and DNA concentration quantification as above. Up to 96 libraries with unique indices were pooled in equimolar concentrations. Library quality was validated on an Agilent TapeStation and concentration was measured via Qubit 2.0 fluorometer and qPCR.

### Single cell sequencing and data pre-processing

Paired-end bulk RNA sequencing (2x150bp) was performed on an Illumina HiSeq sequencer and demultiplexed using Illumina bcl2fastq (v2.20). Reads were trimmed of adaptor sequences, quality filtered, and de-duplicated in fastp (v0.23.2) and aligned to reference sequence hg19 with bowtie2 (v2.3.5). Format conversions were performed with samtools (v1.9) and bedtools (v2.29.2).

### Single cell copy number calling

Copy number calls were performed in a local installation of Ginkgo (69) with a variable bin size of ∼2.5 Mb using global segmentation. Minimum ploidy was set to 1.35 and maximum ploidy to 3 to reflect our flow cytometric gating strategy for FACS. Whole chromosome copy numbers were estimated by taking the mode of copy numbers across genomic bins for each chromosome. Thus, even sub-chromosomal gains and losses of greater than 50% of a chromosome should remain detectable.

### Quantification of single cell CIN signatures

We quantified previously published putative CIN signatures from Drews et al. 2022 using resources made publicly available in a github repository by the authors (https://web.archive.org/web/20220615195321/https://github.com/markowetzlab/CINSignatureQuantification). We used the function ‘quantifyCNSignatures’ in the available R package CINSignatureQuantification (44) to call signatures from unrounded segment copy number matrices derived using Ginkgo, as described above and report the normalized, unscaled signature activity levels to avoid masking the relative activity of each signature.

### Inference of mis-segregation rates

We used agent-based simulation and approximate Bayesian computation to infer mis-segregation rates from scDNAseq data. Agent-based simulation of CIN and karyotype selection in growing populations was performed in NetLogo (v6.0.4) (70) and approximate Bayesian computation was performed using the R (v4.2.2) (67) using the abc package (v2.1)(71) as previously described (27).

#### Agent based simulation

We initiated simulated populations with 100 diploid cells and evolved exponentially growing populations using a pseudo-Moran process to reduce computational demands (a random 50% of cells are culled when the population surpasses 3000 cells). Euploid cells had a 50% chance to divide at every step, a probability that is modified according to a cell’s fitness level. We used a karyotype selection scheme that emulates stabilizing selection by negatively selecting genetically unbalanced karyotypes. The contribution of each chromosome to karyotype selection is dependent on the abundance of genes it encodes. We simulated populations using the following parameters: MDD = [0, 0.046, 0.092, …, 46], S = [0, 1, 2, …, 200], Time Steps = [0, 1, 2, …, 100]. We assumed whole-chromosome mis-segregation and that chromosome copy numbers below 1 and above 7 would result in cell death. Accordingly, as cells divide and mis-segregate chromosomes, more aneuploid cells with more unbalanced karyotypic stoichiometries are less likely to continue division. A random selection of 300 karyotypes is exported to measure summary statistics at each time step. Each combination was repeated 3 times for a total of 1,006,005 independent simulations. See Lynch et al. 2022 for additional details.

#### Population summary statistics

To summarize the characteristics of karyotypes from simulated populations and scDNAseq data, we quantified three features: aneuploidy, mean karyotype variance (MKV), and Colless index. Aneuploidy was quantified as the mean variance of copy numbers within each cell’s sub-clonal karyotype (normalized to the modal karyotype). MKV was quantified as the mean variance of copy numbers for each chromosome across the population. Colless index was measured using the R package phyloTop (v2.1.1)(72) and was normalized to the number of leaf tips. Phylogenetic trees for measuring the Colless index were reconstructed from chromosome copy numbers by computing Euclidean distance matrices and hierarchical clustering using complete linkage, both in the R stats package (v.4.2.3)(67). Hierarchical clustering may not produce the same results every time, and rare, highly different dis-similar observations can drastically skew Colless indices, both of which result in measurement noise. To reduce this noise we, we permute the copy numbers of individual homologous chromosomes (i.e., a population’s copy numbers for chromosome 1 are permuted separately from those of chromosome 2) across the population and repeat this Colless index measurement, taking the average of 200 iterations. This preserves phylogenetic imbalance for populations with sub-clonal alterations but reduces imbalance for those with a rare highly dissimilar karyotype, resulting in a stable and reproducible measure. Also, because hierarchical clustering requires at least 3 observations, we removed all simulated datasets that failed to maintain at least 3 cells, which could occur when high mis-segregation rates force cells past the pre-defined lethal chromosome copy number thresholds (1 to 7), promoting population collapse.

#### Approximate Bayesian computation

Parameter inference of mis-segregation rates and selection pressure from scDNAseq data was performed with approximate Bayesian computation using our simulated dataset as a prior dataset. We used the R package abc (v2.1) (71) to complete the analysis with rejection sampling and a tolerance threshold of 0.01. We chose independent prior datasets that best matched each control and experimental group. For example, we assigned control groups a prior dataset that encompassed 30-50 time steps (i.e., 15-25 generations) to reflect the number of passages after mono-clonal culture. Groups with doxycycline-induced CIN were assigned a prior dataset with ≤ 4 time steps (2 generations) to encompass about 48 hours of growth with fully penetrant CIN induction (this assumes induction of the tetOn constructs takes about 24 hours to become fully induced). The polar chromosome model (Po), which was induced using sequential chemical treatments and enrichment of about 50% of the population, was assigned a prior dataset with ≤ 2 time steps (1 generation).

### Bulk DNA sequencing and analysis

#### Bulk sample and library preparation

Sample preparation, sorting, and bulk DNA library preparation were prepared in parallel with and in the same manner as single-cell DNA samples. 500 cells were sorted for each bulk DNA sample.

#### Bulk DNA sequencing and data pre-processing

Paired-end bulk DNA sequencing (2x150bp) was performed on an Illumina HiSeq to ∼10x coverage and demultiplexed using Illumina bcl2fastq (v2.20). Sequencing reads were adaptor-trimmed, filtered, de-duplicated, and aligned to hg19 with the Illumina DRAGEN Bio-IT Platform using default settings.

#### Bulk copy number calling

Bulk DNA copy numbers were called in R using QDNAseq (v1.34.0) (73) and a bin size of 30 Kb. Segment copy numbers were called using bin copy numbers smoothed over 2 bins and Anscombe transformed (transformFun = “sqrt” in the ‘segmentBins’ function).

#### Quantification of CIN signatures

Putative CIN signatures were derived from bulk copy number profiles and reported in the same manner as for single cell DNA copy number profiles as described above.

### Bulk RNA sequencing

#### Bulk RNA sample preparation

Cells were re-seeded in 6-well plates at 40% and grown to ∼70-80% density over 18-24 hours prior to harvest. Media was removed from cells and immediately lysed with TRI Reagent (ThermoFisher Scientific, AM9738) to preserve RNA, which was stored at −80°C. Total RNA was isolated using Zymo Direct-zol RNA MiniPrep kit (Zymo Research, R2050) and the concentration and quality were assessed with a Qubit 2.0 fluorometer and Agilent TapeStation respectively.

#### Bulk RNA library preparation

RNA libraries were prepared using the NEBNext Ultra II RNA Illumina Library Prep Kit (New England Biolabs, E7775) and mRNAs were enriched with oligo-d(T) beads then fragmented for 15 minutes at 94°C prior to first and second strand cDNA synthesis. cDNAs were end-repaired, 3’-adenylated, ligated with universal Illumina adapters and unique index sequences, then enriched by PCR. Library quality was validated on an Agilent TapeStation and concentration was measured via Qubit 2.0 fluorometer and qPCR.

#### Bulk RNA sequencing, data pre-processing, and analysis

Paired-end bulk RNA sequencing (2x150bp) was performed on an Illumina HiSeq sequencer and demultiplexed using Illumina bcl2fastq (v2.20). Demultiplexed reads were trimmed of universal Illumina index sequences with BBMap’s bbduk (v38.95) (74). We aligned trimmed reads to reference sequence hg19 and generated fragment count matrices in R/Rstudio (v4.2.3/v1.2.5019) (67, 75) using Rsubread (v2.4.3) (76). Library size normalization and dispersion estimation was performed in edgeR (v.3.32.1) (77) while linear modeling and differential expression analysis was performed in limma (v3.46) (78, 79).

## AUTHOR CONTRIBUTIONS

Conceptualization: ARL, BAW, and MEB.

Methodology: ARL, ASZ, KO, LH.

Data collection: ARL, KO, LH.

Formal analysis: ARL, SB.

Data curation: ARL.

Writing – original draft: ARL.

Writing – review and editing: ARL, SB, ASZ, KO, LH, VLH, BAW, and MEB.

Data visualization: ARL.

Supervision, project administration, and funding acquisition: VLH, BAW, and MEB.

### ACKNOWLEDGMENTS

We thank the University of Wisconsin Carbone Cancer Center (UWCCC) Shared Resources funded by the UWCCC Support Grant P30 CA014520 – Flow Cytometry Core Facility (1S10RR025483-01), Cancer Informatics Shared Resource, Small Molecule Screening Facility. Thanks also go to the Wisconsin State Laboratory of Hygiene for cytogenetic expertise and support, Dr. Rob Lera for microscopy assistance, and members of the laboratories of Drs. Mark Burkard, Beth Weaver and Aussie Suzuki for many productive discussions.

## COMPETING INTEREST STATEMENT

MEB declares all interests without adjudicating relationship to the published work. He is on the medical advisory board of Strata Oncology, receives research funding from Abbvie, Genentech, Puma, Arcus, Apollomics, Loxo Oncology/Lilly, and Elevation Oncology, and holds patents on a microfluidic device for drug testing and for homologous recombination and super-resolution microscopy technologies.

All other authors report no conflicts of interest.

## CODE AND DATA AVAILABILITY

Scripts and data used for data analysis and visualization will be freely available on Open Science Framework and sequence data will be deposited to NCBI SRA and GEO prior to final publication.

## FINANCIAL SUPPORT

This work was supported by NIH grants 1R01CA234904 (MEB, BAW) and P30CA014520 (UWCCC Support Grant). ARL was supported by F31CA254247 and ASZ was supported by T32GM008688.

---

**Supplemental Table 1.**
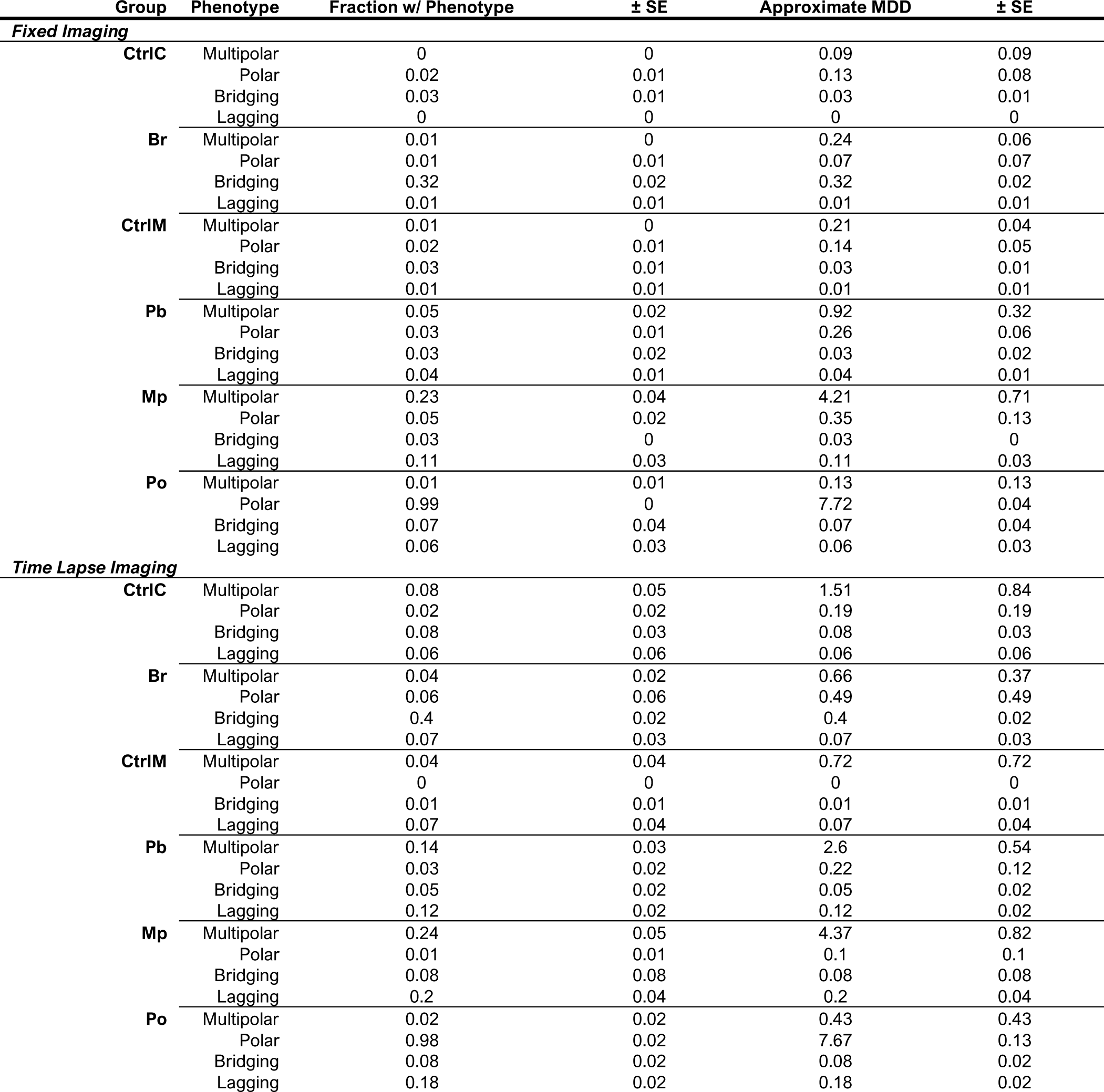
**Approximated MDD by phenotype** Approximated MDD for each CIN phenotype in each model. MDD was calculated using 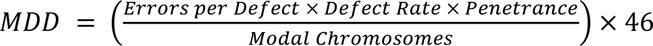 and assumptions of the number of chromosomes mis-segregated for each defect (see Materials and Methods). The fraction of cells with polar chromosomes represents metaphase cells wherein they are most readily detectable. All other CIN phenotypes are taken from anaphase or telophase cells.

| Group | Phenotype | Fraction w/ Phenotype | ± SE | Approximate MDD | ± SE |
| --- | --- | --- | --- | --- | --- |
| *Fixed Imaging* |  |  |  |  |  |
| **CtrlC** | Multipolar | 0 | 0 | 0.09 | 0.09 |
|  | Polar | 0.02 | 0.01 | 0.13 | 0.08 |
|  | Bridging | 0.03 | 0.01 | 0.03 | 0.01 |
|  | Lagging | 0 | 0 | 0 | 0 |
| **Br** | Multipolar | 0.01 | 0 | 0.24 | 0.06 |
|  | Polar | 0.01 | 0.01 | 0.07 | 0.07 |
|  | Bridging | 0.32 | 0.02 | 0.32 | 0.02 |
|  | Lagging | 0.01 | 0.01 | 0.01 | 0.01 |
| **CtrlIM** | Multipolar | 0.01 | 0 | 0.21 | 0.04 |
|  | Polar | 0.02 | 0.01 | 0.14 | 0.05 |
|  | Bridging | 0.03 | 0.01 | 0.03 | 0.01 |
|  | Lagging | 0.01 | 0.01 | 0.01 | 0.01 |
| **Pb** | Multipolar | 0.05 | 0.02 | 0.92 | 0.32 |
|  | Polar | 0.03 | 0.01 | 0.26 | 0.06 |
|  | Bridging | 0.03 | 0.02 | 0.03 | 0.02 |
|  | Lagging | 0.04 | 0.01 | 0.04 | 0.01 |
| **Mp** | Multipolar | 0.23 | 0.04 | 4.21 | 0.71 |
|  | Polar | 0.05 | 0.02 | 0.35 | 0.13 |
|  | Bridging | 0.03 | 0 | 0.03 | 0 |
|  | Lagging | 0.11 | 0.03 | 0.11 | 0.03 |
| **Po** | Multipolar | 0.01 | 0.01 | 0.13 | 0.13 |
|  | Polar | 0.99 | 0 | 7.72 | 0.04 |
|  | Bridging | 0.07 | 0.04 | 0.07 | 0.04 |
|  | Lagging | 0.06 | 0.03 | 0.06 | 0.03 |
| *Time Lapse Imaging* |  |  |  |  |  |
| **CtrlC** | Multipolar | 0.08 | 0.05 | 1.51 | 0.84 |
|  | Polar | 0.02 | 0.02 | 0.19 | 0.19 |
|  | Bridging | 0.08 | 0.03 | 0.08 | 0.03 |
|  | Lagging | 0.06 | 0.06 | 0.06 | 0.06 |
| **Br** | Multipolar | 0.04 | 0.02 | 0.66 | 0.37 |
|  | Polar | 0.06 | 0.06 | 0.49 | 0.49 |
|  | Bridging | 0.4 | 0.02 | 0.4 | 0.02 |
|  | Lagging | 0.07 | 0.03 | 0.07 | 0.03 |
| **CtrlIM** | Multipolar | 0.04 | 0.04 | 0.72 | 0.72 |
|  | Polar | 0 | 0 | 0 | 0 |
|  | Bridging | 0.01 | 0.01 | 0.01 | 0.01 |
|  | Lagging | 0.07 | 0.04 | 0.07 | 0.04 |
| **Pb** | Multipolar | 0.14 | 0.03 | 2.6 | 0.54 |
|  | Polar | 0.03 | 0.02 | 0.22 | 0.12 |
|  | Bridging | 0.05 | 0.02 | 0.05 | 0.02 |
|  | Lagging | 0.12 | 0.02 | 0.12 | 0.02 |
| **Mp** | Multipolar | 0.24 | 0.05 | 4.37 | 0.82 |
|  | Polar | 0.01 | 0.01 | 0.1 | 0.1 |
|  | Bridging | 0.08 | 0.08 | 0.08 | 0.08 |
|  | Lagging | 0.2 | 0.04 | 0.2 | 0.04 |
| **Po** | Multipolar | 0.02 | 0.02 | 0.43 | 0.43 |
|  | Polar | 0.98 | 0.02 | 7.67 | 0.13 |
|  | Bridging | 0.08 | 0.02 | 0.08 | 0.02 |
|  | Lagging | 0.18 | 0.02 | 0.18 | 0.02 |

**Supplemental Table 2.** **CIN70 and HET70 genes** Genes included in the CIN70 and HET70 gene expression panels. Genes are sorted in their original, published order, CIN70 by correlational rank and HET70 in alphabetical order.

| CIN70 |  | HET70 |  |
| --- | --- | --- | --- |
| TPX2 | MSH6 | AHCYL1 | LPP |
| PRC1 | EZH2 | AKT3 | MED8 |
| FOXM1 | CTPS1 | ANO10 | MMP2 |
| CDK1 | DKC1 | ANTXR1 | MUL1 |
| TGIF2 | OIP5 | ATP6V0E1 | MYO10 |
| MCM2 | CDCA8 | ATXN1 | NAGK |
| H2AZ1 | PTTG1 | B4GALT2 | NR1D2 |
| TOP2A | CEP55 | BASP1 | NRIP3 |
| PCNA | H2AX | BHLHE40 | P4HA2 |
| UBE2C | CMAS | BLVRA | PKIG |
| MELK | NCAPH | CALU | PLOD2 |
| TRIP13 | MCM10 | CAP1 | PMP22 |
| NCAPD2 | LSM4 | CAST | POFUT2 |
| MCM7 | NCAPG2 | CAV1 | POMGNT1 |
| RNASEH2A | ASF1B | CLIC4 | PRKAR2A |
| RAD51AP1 | ZWINT | CTSL | MOK |
| KIF20A | PBK | CYB5R3 | RHOC |
| CDC45 | ZWILCH | ELOVL1 | RRAGC |
| MAD2L1 | CDCA3 | EMP3 | SEC22B |
| ESPL1 | ECT2 | FKBP14 | SERPINB8 |
| CCNB2 | CDC6 | FN1 | SPAG9 |
| FEN1 | UNG | FST | SQSTM1 |
| TTK | MTCH2 | GNA12 | TIMP2 |
| CCT5 | RAD21 | GOLT1B | EMC3 |
| RFC4 | ACTL6A | HECTD3 | TRIM16 |
| ATAD2 | GPI | HEG1 | TRIO |
| CKAP5 | SRSF2 | HOMER3 | TUBB2A |
| NUP205 | HDGF | IGFBP3 | VEGFC |
| CDC20 | NXT1 | IL6ST | VIM |
| CKS2 | NEK2 | ITCH | WASL |
| RRM2 | DHCR7 | P3H1 | YIPF5 |
| ELAVL1 | AURKA | P3H2 | YKT6 |
| CCNB1 | NDUFAB1 | LEPROT | ZBTB38 |
| RRM1 | NEMP1 | LGALS1 | ZCCHC24 |
| AURKB | KIF4A | LIMA1 | ZMPSTE24 |

## SUPPLEMENTAL FIGURE LEGENDS

**Supplemental Figure 1.**
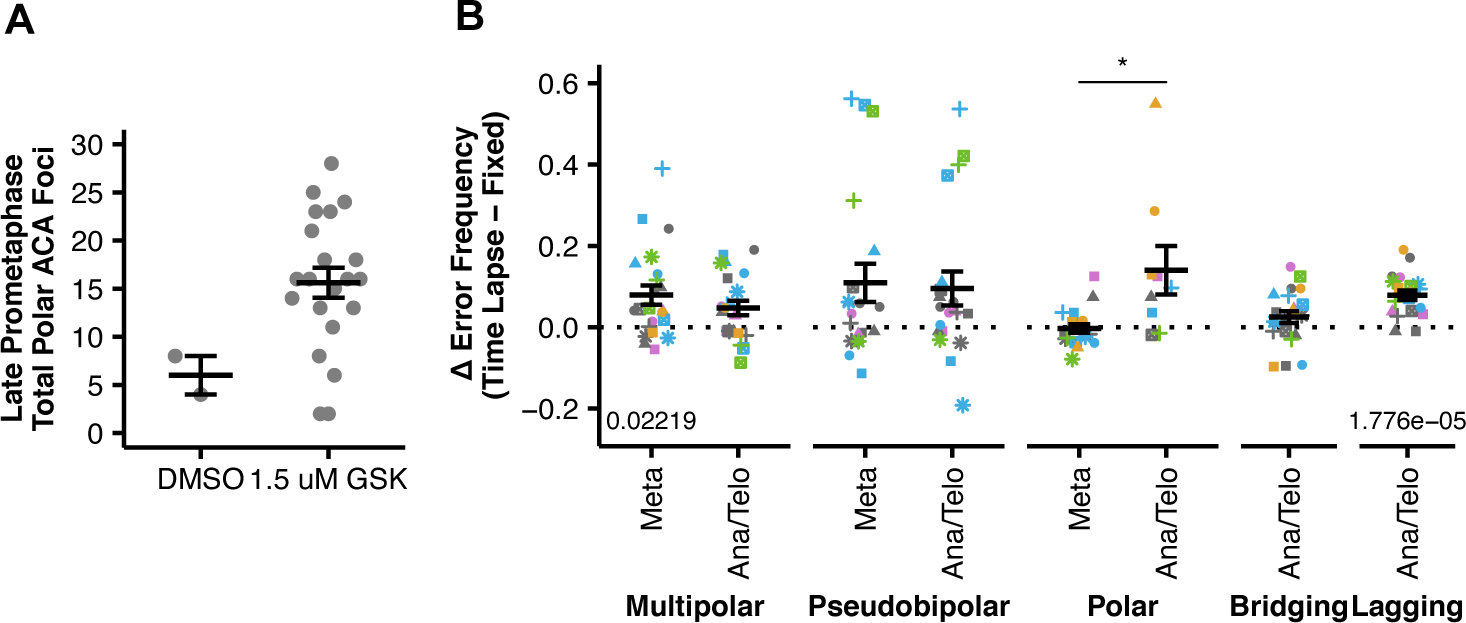
Additional Imaging Data. **(A)** Number of anticentromere antibody (ACA) foci at or behind the spindle pole in late prometaphase (i.e., discernable metaphase plates) MCF10A cells treated under the Po condition but prior to anaphase induction. **(B)** Differences in observed frequencies of mitotic defects between fixed immunofluorescence and time lapse fluorescence imaging in metaphase or anaphase/telophase. Significance values beneath data are from two-tailed, one-sample Student’s *t-*tests where H_o_: μ = 0. Colors of individual points indicate the biological replicate. Bars and error bars indicate mean and standard error. Significance values above data are from a two-tailed, two-sample Student’s *t*-tests.

**Supplemental Figure 2.**
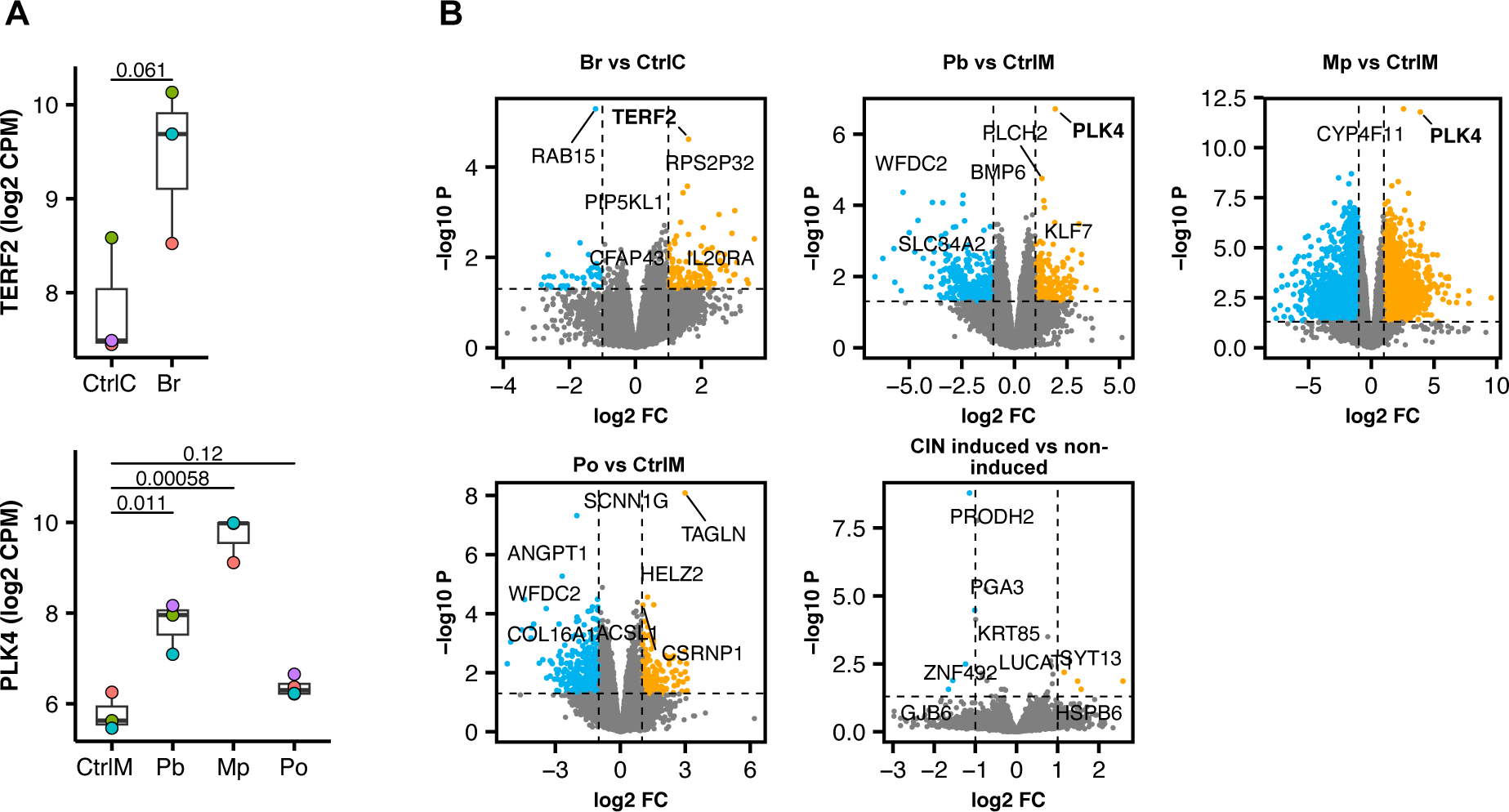
Gene expression in inducible phenotypic models of CIN. **(A)** Expression levels (log2 counts per million (CPM)) of *TERF2* and *PLK4* in CAL51-TERF2-DN-tetOn and MCF10A-PLK4-WT-tetOn cells respectively. Two-tailed, two-sample Student’s *t*-tests are shown above data. N ≥ 3 biological replicates. **(B)** Volcano plots of differential gene expression in each CIN-induced model compared to its uninduced control and a pooled analysis of all CIN-induced groups compared to all non-induced groups. Dashed lines indicate log2 fold-change thresholds of -1/1 and unadjusted P value of 0.05.

**Supplemental Figure 3.**
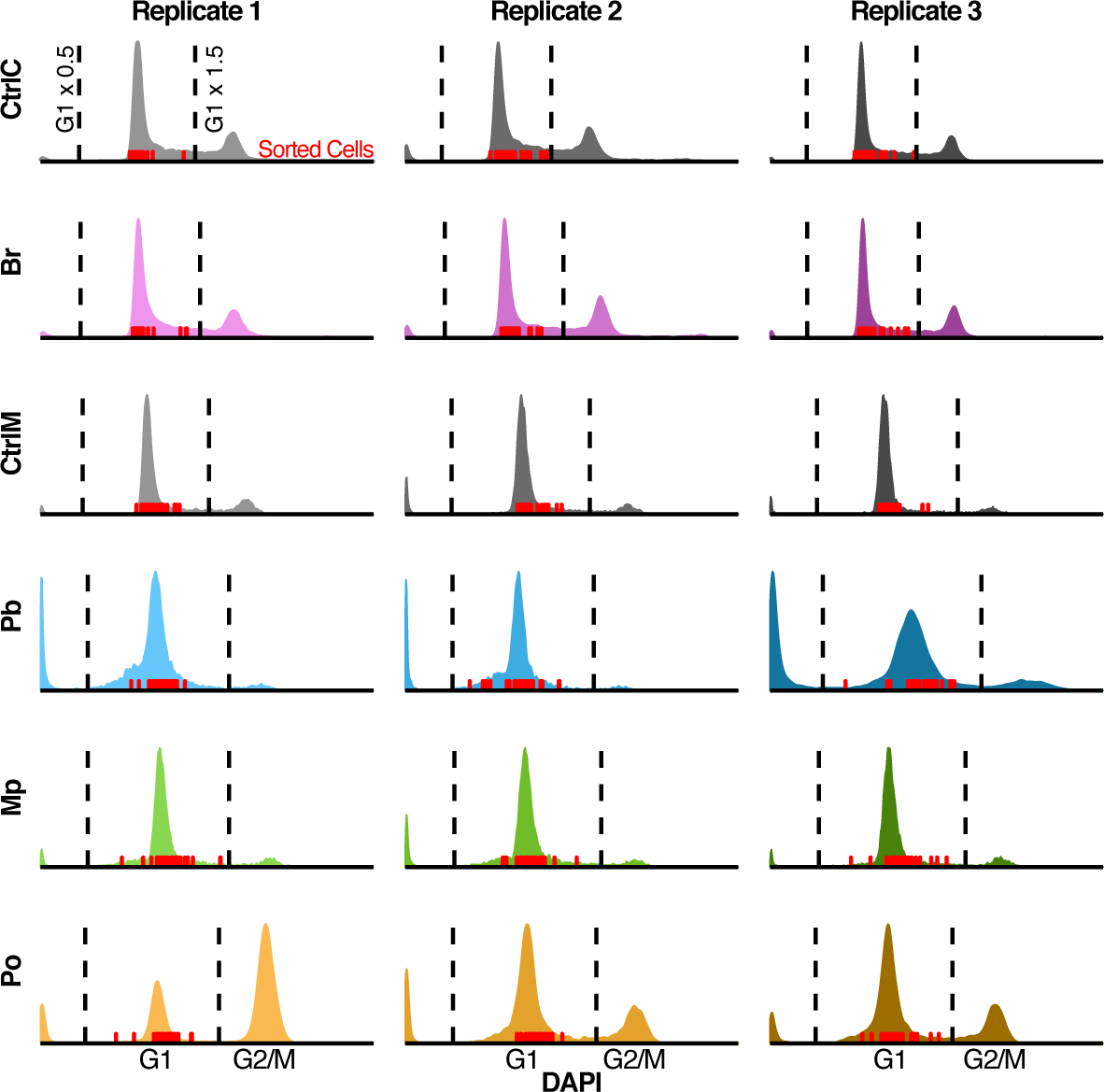
DNA content analysis and gating for pre-scDNAseq FACS. Cell cycle profiles (measured by flow cytometric analysis of DAPI intensity) for each replicate of each phenotypic model of CIN. Dashed lines indicate gating strategy for FACS of cells for scDNAseq and were determined by 50% and 150% of the intensity of the G1 peak. Red ticks indicate the DAPI intensity of cells sorted for scDNAseq. Note the broader G1 peaks of CIN-induced models Pb, Mp, and Po, indicative of extensive aneuploidy.

**Supplemental Figure 4.**
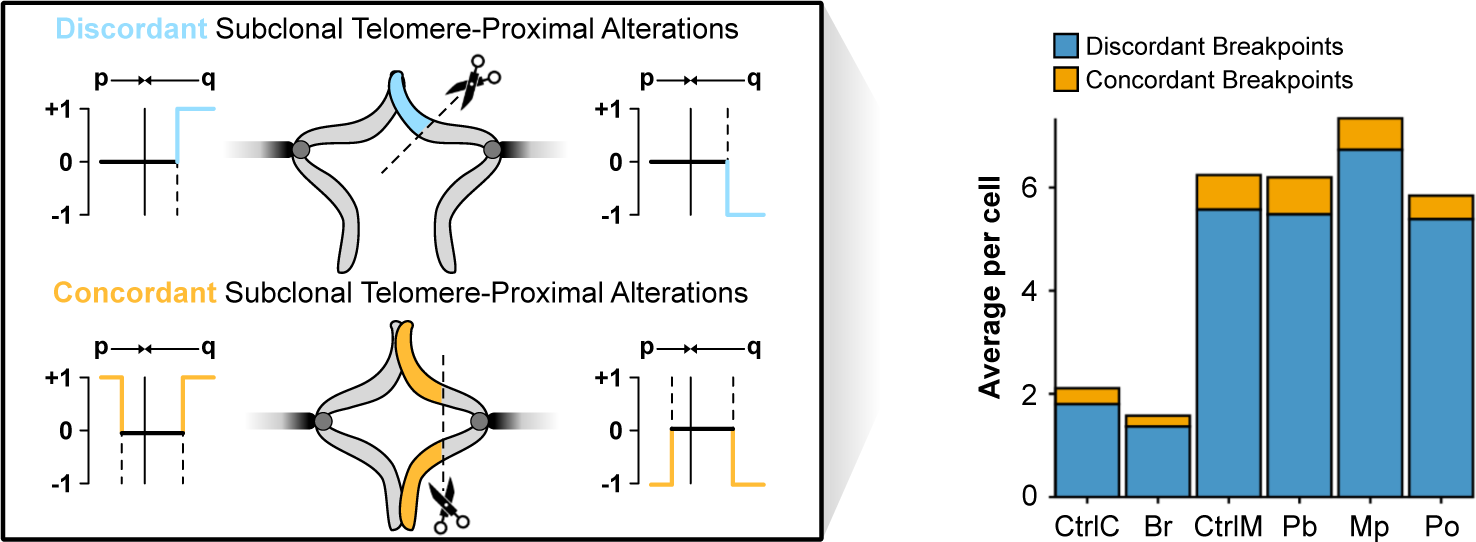
Breakpoint analysis in scDNAseq data. Breakpoint analysis of discordant and concordant telomere-proximal copy number alterations. Copy numbers are normalized relative to those of the modal karyotype of each phenotypic CIN model to uncover subclonal or relatively recent alterations. Only segments with copy number alteration consensus across the 4 telomere-proximal genomic bins (10 Mb), but which terminate at or before the centromere, are considered. Discordant subclonal telomere-proximal alterations are defined as those that are not matched by an alteration on the opposite chromosome arm. Concordant alterations are those that are matched on the opposite chromosome arm. The quantification shows the average number of each alteration per cell for each model.

**Supplemental Figure 5.**
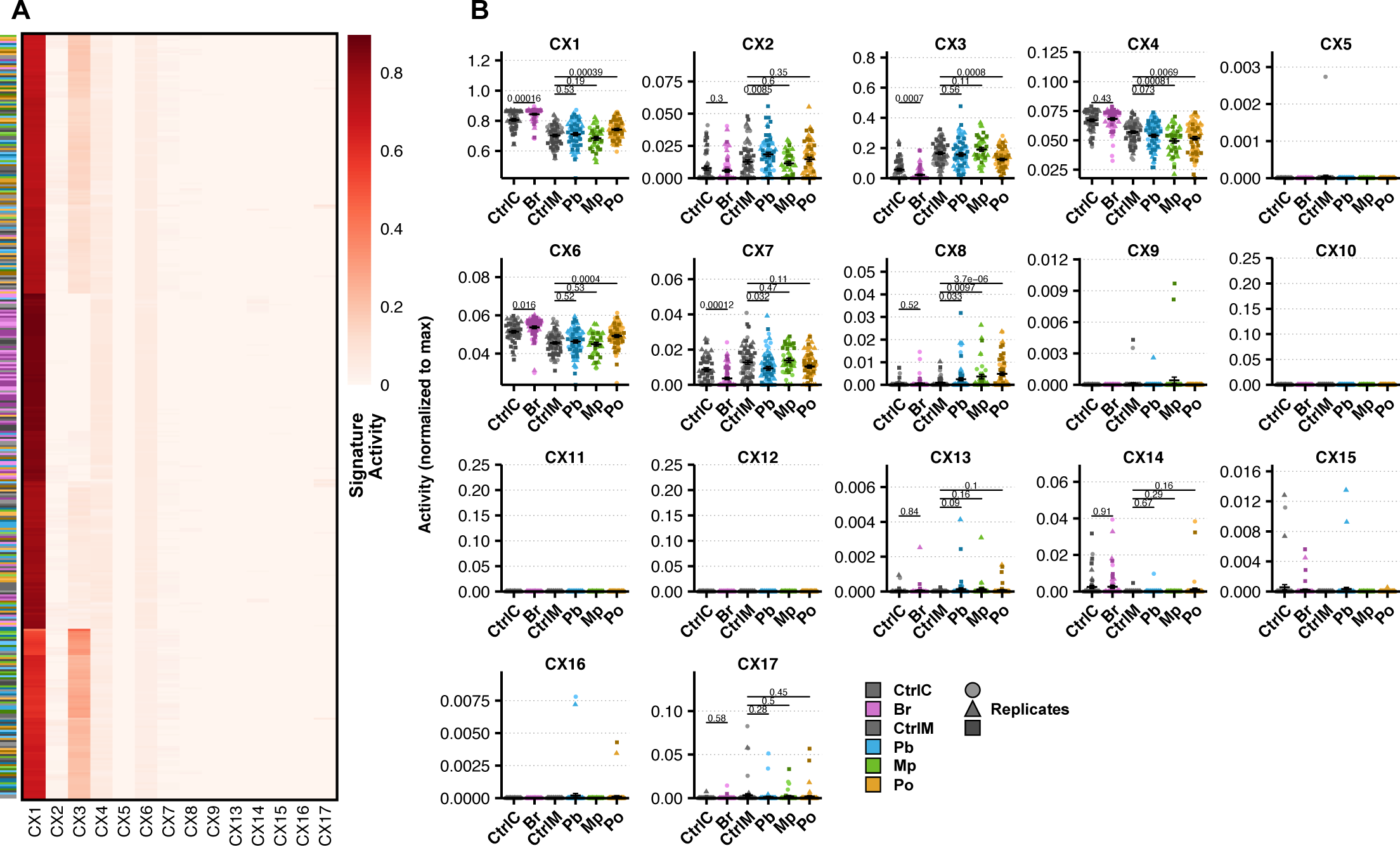
Putative CIN signatures in inducible phenotypic models of CIN at single cell resolution. **(A)** Heatmap of all normalized putative CIN signature measurements in single cells showing with rows clustered by similarity. Colorized annotation on the left indicates phenotypic CIN model and shade indicates biological replicate. **(B)** Normalized putative CIN signature measurements in single cells grouped by model. Results from two-tailed, two-sample Student’s *t*-tests are shown above data using all cells across 3 biological replicates. Shape and shade of data points indicate biological replicate.

**Supplemental Figure 6.**
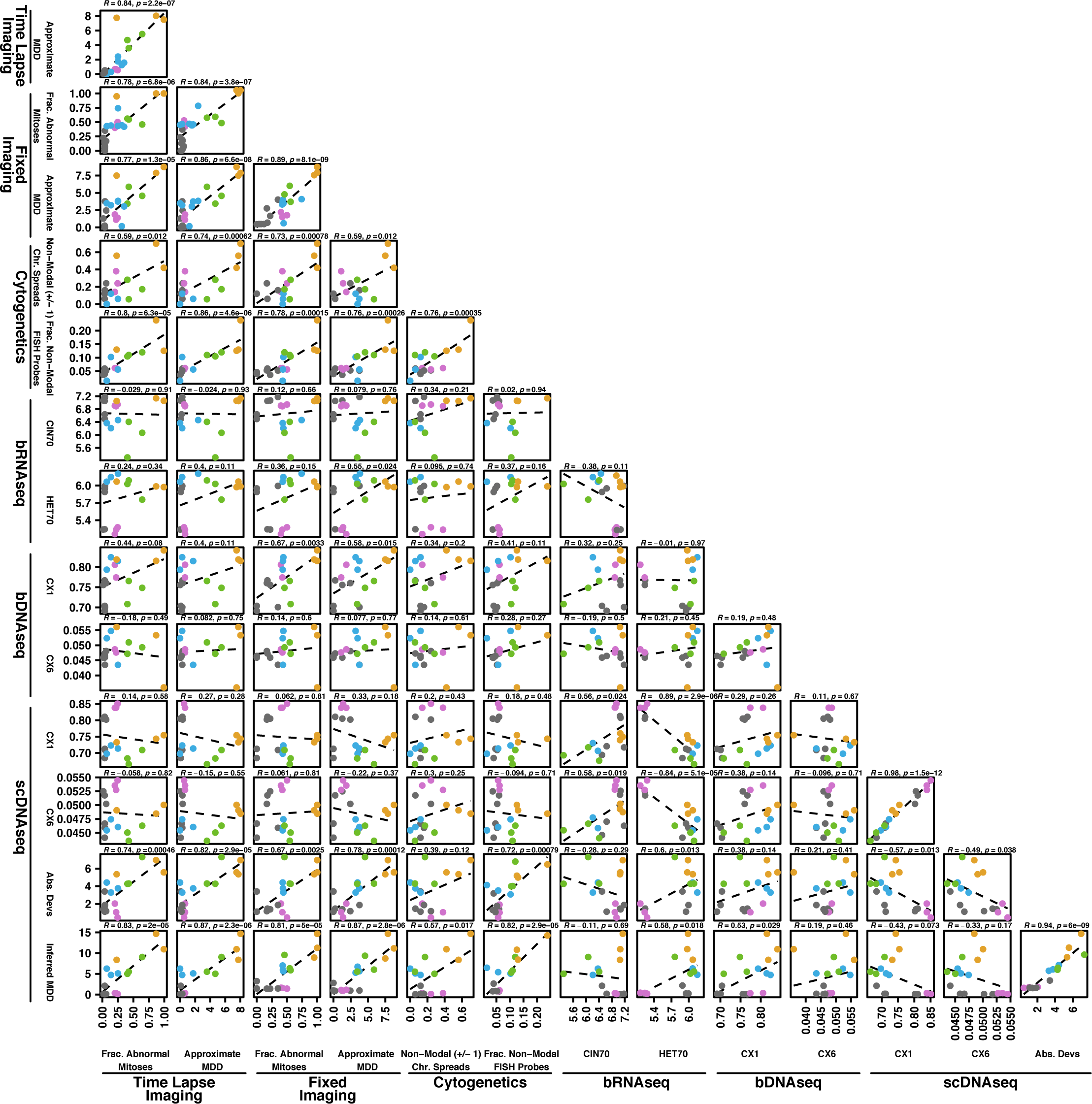
Extended pairwise correlations of CIN measurements. Pairwise correlations between each CIN measurement. Colors of data points indicate the phenotypic model of CIN. Pearson correlation coefficients and p-values of regression are shown.

## Notes

### Competing Interest Statement

MEB is on the medical advisory board of Strata Oncology, receives research funding from Abbvie, Genentech, Puma, Arcus, Apollomics, Loxo Oncology/Lilly, and Elevation Oncology, and holds patents on a microfluidic device for drug testing and for homologous recombination and super-resolution microscopy technologies.
All other authors report no conflicts of interest.

